# Effects of Oncohistone Mutations and PTM Crosstalk on the N-terminal Acetylation Activities of NatD

**DOI:** 10.1101/2021.10.21.465300

**Authors:** Yi-Hsun Ho, Rong Huang

## Abstract

Acetylation at the α-N-terminus (Nα) is one of the most abundant modifications detected on histone H4 and H2A, which is catalyzed by N-terminal acetyltransferase D (NatD). NatD substrate recognition motif, N-terminal SGRGK, is enriched with frequent oncohistone mutations and post-translational modifications (PTMs). However, there is no information on how oncohistone mutations and other PTMs affect NatD-catalyzed acetylation. Herein, we determined how changes of local chemical environment on the N-terminal SGRGK sequence regulate NatD-catalyzed Nα acetylation on histone H4/H2A. Our studies indicate that all oncohistone mutations at SGRG suppressed the catalytic efficiency of NatD. Meanwhile, H4 serine 1 phosphorylation and arginine 3 methylation also negatively affect the NatD activity, but the lysine 5 acetylation has a marginal effect on NatD. This work reveals the impacts of oncohistone mutations on NatD activity and unravels the crosstalk between NatD and PTMs. To our knowledge, this is the first report on the potential regulatory mechanism of NatD, highlighting different revenues to interrogate the NatD-mediated pathway in the future.

## Introduction

Histone modifications alter the chromatin structures and DNA accessibility, serving as a critical mechanism in the regulation of gene expression, DNA replication, and DNA repair.^1,2^ Concerning the modification type and site, at least 450 modifications have been identified on histones, such as phosphorylation, methylation, and acetylation.^3^ Crosstalk among these modifications adds another dimension to the dynamical regulation of transcription.^2^ The α-N-terminal (Nα) acetylation catalyzed by N-terminal acetyltransferases (NATs) is a predominantly modification for approximate 80% of proteins in humans.^4–6^ NATs have been classified into NatA– H based on their subunit compositions and substrate specificity profiles.^6^ Among NATs, NatD specifically acetylates an SGRG sequence at the protein N-terminus (Figure 1A). Until now, H4 and H2A are the only two confirmed protein substrates for NatD.^7–10^ Notably, Nα-acetylation of H4 (Nα-acH4) and H2A (Nα-acH2A) were the most abundant marks of histone proteins through several proteomic investigations.^11–16^ For instance, over 80% of H4 and H2A were found Nα-acetylated in mouse brains.^14^ In addition, 97–98% of Nα-acH4 was reported in breast cancer cell lines.^11^

**Figure 1.**
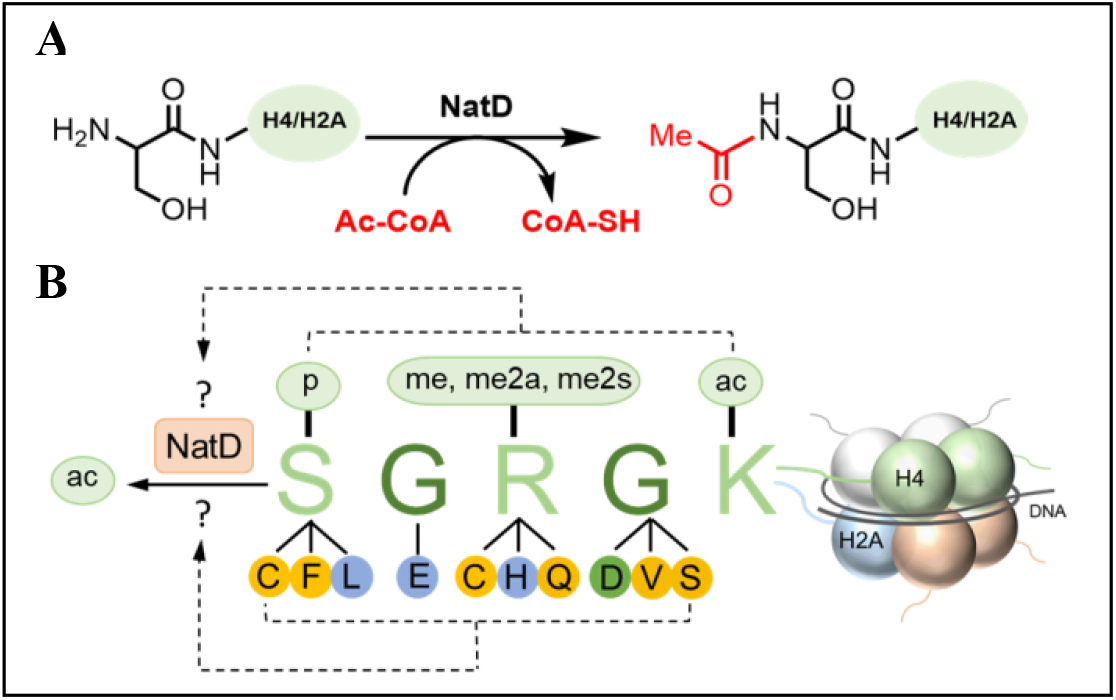
(A) Nα-acetylation catalyzed by NatD. (B) Mutations of oncohistone H4 and H2A and chemical modifications at residues Ser-1, Arg-3, and Lys-5 on the H4 substrate. Modifications include phosphorylation (p), methylation (me), and acetylation (ac). Mutations detected on H4 and H2A: yellow circle; mutations detected only on H4: green circle; mutations detected only on H2A: blue circle.

Recently, oncohistones have drawn significant attention because of their oncogenic features and recurrence at the epigenetic modification-rich histone tails.^17–21^ Interestingly, multiple mutations have been identified in all four amino acids of the SGRG recognition motif of NatD (Figure 1B).^21–24^ Notably, Ser-1 and Arg-3 are the top two frequently mutated residues of H4, and Ser-1 is the most frequently mutated residue of H2A.^23^ Although the detailed mechanism of these mutations remains enigmatic, it raises the question of how those oncohistone mutations at the N-terminal regions affect NatD-catalyzed Nα-acetylation.

Meanwhile, the H4 tail is abundant with post-translational modifications (PTMs), including phosphorylation, methylation, and acetylation. Nα-acH4 has been found to affect other PTMs at Ser-1, Arg-3, and Lys-5 on H4 to influence biological outcomes.^25–30^ For instance, Nα-acH4 interferes with asymmetrical dimethylation on H4 Arg-3 residue (H4R3me2a) in yeast.^25,26^ Caloric restriction in yeast suppresses NatD expression and increases the ratio of H4R3me2a to Nα-acH4, promoting the expression of metabolic and stress-response genes.^26^ Consistent with the findings in yeast, a biochemical study demonstrated that Nα-acetylation on H4 peptide reduced the rate of H4R3me2a catalyzed by protein arginine methyltransferase 1 (PRMT1) and PRMT3 but had a marginal effect on the Arg-3 dimethylation by PRMT5 and PRMT8.^28,30^ In colorectal cancer, loss of N-acH4 decreased symmetric dimethylation on H4 Arg-3 (H4R3me2s) levels by downregulating PRMT5, but has marginal effect on monomethylation (H4R3me1) or H4R3me2a.^29^ Nα-acH4 has been demonstrated to block casein kinase 2α-mediated phosphorylation at the Ser-1 (H4pS1) in lung cancer cells, inducing Slug expression and metastasis.^27^ Additionally, loss of NatD alters the levels of H4R3me2s, H4R3me1, H4R3me2a, and acetylation of H4 Lys-5 (H4K5ac).^29^ Although Nα-acetylation on H4 has exhibited to impact Arg-3 methylation, Ser-1 phosphorylation, and Lys-5 acetylation, it remains elusive if the communication is unidirectional or mutual. Because the first four residues SGRG are critical for substrate recognition, we hypothesize that modifications on SGRG may affect NatD-catalyzed Nα-acetylation. Herein, we examine how the oncohistone mutations and post-translational modifications of H4S1, H4R3, and H4K5 affect Nα-acetylation by NatD (Figure 1B).

## Results and Discussion

### Effect of histone H4 length on NatD activity

Previous studies suggested that yeast NatD requires a more extended substrate peptide sequence for Nα-acetylation than human NatD (hNatD).^7^ Specifically, complete acetylation was detected in yeast when the first 50 residues of H4 were present but incomplete in the presence of the first 30 residues.^7^ In contrast, hNatD displayed a comparable catalytic efficiency for H4 synthetic peptides consisting of the first 5, 8, and 19 residues in a biochemical radioactive study.^10^ To systematically understand how substrate length affects hNatD catalysis, we determined the steady-state kinetic parameters for both full-length H4 protein and a panel of truncated H4 synthetic peptides using recombinant hNatD and a continuous fluorescence assay.^31^ The Michaelis-Menten parameters are comparable for H4 protein and the cofactor acetyl coenzyme A (AcCoA) (Table 1). Indeed, hNatD exhibited comparable catalytic activity (*k*_cat_) for its substrates with variable lengths ranging from full-length protein to a tripeptide (Figure 2). However, the catalytic efficiency (*k*_cat_/*K*_m_) of hNatD altered for different lengths of substrates, which is mainly due to decreased binding affinity as reflected by *K*_m_ values.

**Table 1.** Effect of C-terminal truncations of hNatD recognition.

| ID | Sequence | $K_m$ ($\mu\text{M}$) | $k_{\text{cat}}$ ($\text{min}^{-1}$) | $k_{\text{cat}}/K_m$ ($\text{M}^{-1} \text{min}^{-1}$) |
| --- | --- | --- | --- | --- |
| Full length Histone H4 | | $0.80 \pm 0.10$ | $12 \pm 0.42$ | $1.5 \times 10^7$ |
| H4-21 | SGRGKGGKGLGKGGAKRHRKV | $0.83 \pm 0.095$ | $11 \pm 0.37$ | $1.3 \times 10^7$ |
| H4-12 | SGRGKGGKGLGK | $2.1 \pm 0.22$ | $11 \pm 0.35$ | $5.2 \times 10^6$ |
| H4-8 | SGRGKGGK | $3.8 \pm 0.38$ | $10 \pm 0.26$ | $2.6 \times 10^6$ |
| H4/H2A-5 | SGRGK | $5.9 \pm 0.59$ | $11 \pm 0.46$ | $1.9 \times 10^6$ |
| H4/H2A-4 | SGRG | $28 \pm 2.4$ | $11 \pm 0.39$ | $3.9 \times 10^5$ |
| H4/H2A-3 | SGR | $211 \pm 21$ | $9.5 \pm 0.48$ | $4.5 \times 10^4$ |
| AcCoA | AcCoA | $0.63 \pm 0.055$ | $9.2 \pm 0.17$ | $1.5 \times 10^7$ |

**Figure 2.**
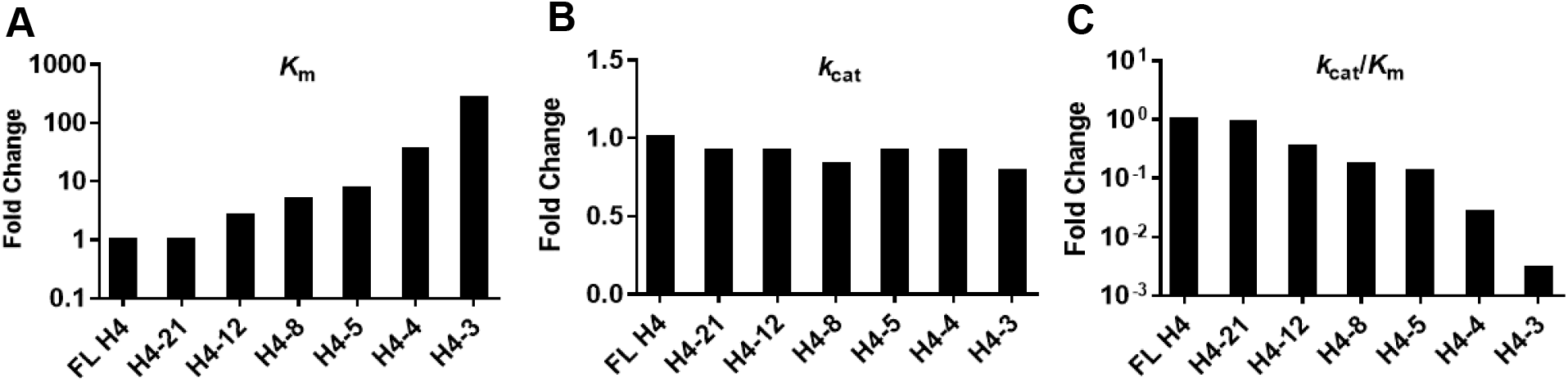
Nα-acetylation of histone H4 peptides with variable length by hNatD. Fold change of (A) *K*_m,_ (B) *k*_cat_, and (C) *k*_cat_/*K*_m_ normalized to full-length histone H4 (FL H4).

Notably, hNatD demonstrated equivalent activity and efficiency for both H4-21 and full-length H4 protein, implying that the C-terminal sequence after Val-21 does not contribute to hNatD recognition. When peptide length decreased from H4-21 to H4-12, no changes were observed for the catalytic activity of hNatD bound with substrate molecules as reflected by *k*_cat_ values (11 ± 0.37 min^-1^ and 11 ± 0.35 min^-1^, respectively). However, there was about a 2.5-fold decrease for H4-12 in catalytical efficiency due to increased *K*_m_ values. Similar trends were observed when peptide length reduced from H4-12 to H4-8 residues, while the kinetic parameters were comparable for H4-8 and H4-5 peptides. When the peptide length decreased from H4-5 to H4-4 and H4-4 to H4-3, the *K*_m_ values increased about 5-fold and 8-fold, respectively. This result agrees with the previous NatD crystal structure, certifying that the first four N-terminal residues are crucial for NatD-specific binding and acetylation.^9,10^

In summary, the *K*_m_ values of six investigated peptide substrates were inversely proportional to their length. The *k*_cat_ values were comparable for peptide substrates with varying lengths from 3–21 residues. These results suggest that the C-terminal truncation (5–20 residues) of H4 substrate has modest effects on the binding towards hNatD but a negligible effect on the catalytic rate of hNatD. Thus, the specificity constant of hNatD is proportional to the peptide length when peptide length is shorter than 21-mer.

### Nα-acetylation of Oncohistone H4 and H2A

The H4 and H2A are the only two validated substrates of hNatD, containing an identical SGRGK motif at their N-termini. The high-frequency mutations observed in the first four residues of H4/H2A prompted us to investigate how Nα-acetylation is affected by H4/H2A oncohistone mutations. Hence, we synthesized pentapeptides derived from the N-terminus of oncohistones H4/H2A and determined their steady-state kinetic parameters using a continuous fluorescence assay (Table 2). Since the fluorescence assay monitors the acetylation progression by detecting the thiol group of CoA products, the existence of cysteine residue in S1C and R3C mutant pentapeptides would interfere with this fluorescence assay. Therefore, we utilized a previously developed MALDI-MS assay to directly detect acetylated products to monitor the acetylation progress of S1C and R3C mutant.^32^ Because the cysteine is prone to form dimers, a reducing agent dithiothreitol (DTT) was added to the reaction mixture to prevent dimer formation. For 1 mM of cysteine-containing peptides, 5 mM DTT was selected to completely suppress the formation of the dimer. To compare the performance of two assays and the effect of DTT, we used H4-5 as control and performed kinetic analysis for H4-5 using the MALDI-MS assay in the presence or absence of 5 mM DTT (Figure S1). The Michaelis-Menten constants obtained from the MALDI-MS assay are comparable to those from the fluorescence assay, validating the feasibility of comparing results from two different assays for those mutant peptides containing a cysteine residue.

**Table 2.** Effects of oncohistone mutations on N-terminal acetylation of H4/H2A peptides

| Peptide ID | H4<br>Frequency | H2A<br>Frequency | Sequence | $K_{\text{m}}$<br>$\mu\text{M}$ | $k_{\text{cat}}$<br>$\text{min}^{-1}$ | $k_{\text{cat}}/K_{\text{m}}$<br>$\text{M}^{-1} \text{ min}^{-1}$ |
| --- | --- | --- | --- | --- | --- | --- |
| H4/H2A-5 | | | SGRGK | $5.9 \pm 0.59$ | $11 \pm 0.46$ | $1.9 \times 10^6$ |
| H4/H2A S1C | High | High | CGRGK | >500 | > 7 | - |
| H4/H2A S1F | Moderate | Low | FGRGK | >5 mM | > 6.8 | - |
| H2A S1L | No | Moderate | LGRGK | $8.8 \pm 1.9$ mM | $2.2 \pm 0.29$ | $2.5 \times 10^2$ |
| H2A G2E | No | Moderate | SERGK | $23 \pm 2.8$ mM | $3.3 \pm 0.31$ | $1.4 \times 10^2$ |
| H4/H2A R3Q | High | High | SGQGK | $393 \pm 35$ | $3.7 \pm 0.12$ | $9.4 \times 10^3$ |
| H2A R3H | Moderate | Low | SGHGK | $26 \pm 1.4$ | $7.5 \pm 0.13$ | $2.9 \times 10^5$ |
| H4/H2A R3C | High | Moderate | SGCGK | 17 | 1.8 | $1.1 \times 10^5$ |
| H4 G4D | High | Low | SGRDK | $272 \pm 40$ | $8.0 \pm 0.66$ | $2.9 \times 10^4$ |
| H4/H2A G4V | Moderate | Low | SGRVK | $37 \pm 1.6$ | $7.7 \pm 0.13$ | $2.1 \times 10^5$ |
| H4/H2A G4S | High | Moderate | SGRSK | $14 \pm 3.1$ | $1.6 \pm 0.16$ | $1.1 \times 10^5$ |

Ser-1 and Gly-2 are two critical residues for substrate-binding with hNatD because the small hydroxyl side chain of Ser-1 and a specific angle of Gly-2 enable the substrate to enter into the substrate-binding pocket of hNatD.^10^ In H4 and H2A, S1C is one of the high-frequency mutations detected in several cancers such as bladder, colorectal, lung, and liver cancers.^21^ We determined the steady-state kinetic parameters of S1C with the highest concentration at 1 mM in the presence of 5 mM of DTT to completely repress its dimer form. We also employed the MALDI-MS assay to determine the kinetics of S1F because of unknown interference from S1F with the fluorescence assay. As shown in Table 2 and Figure 2, hNatD displayed an approximate *k*_cat_ of 7 min^-1^ and 80-fold reduction in *K*_m_ (>500 μM) for S1C mutant that only replaces the hydroxyl side chain of Ser-1 with a thiol group. When the bulky phenylalanine (F) is at the first position, hNatD exhibited a *k*_cat_ of 6.8 min^-1^ and an 800-fold decrease in *K*_m_. For S1L and G2E mutants, the catalytic efficiency was dramatically reduced by over 10,000-fold (*k*_cat_/*K*_m_ = 2.5 × 10^2^ and 1.4 × 10^2^ M^-1^ min^-1^, respectively). These losses are mainly caused by a 1,400 to 3,800-fold reduction in *K*_m_ (S1L *K*_m_ = 8.8 mM and G2E *K*_m_ = 23 mM, respectively), while their catalytic rates *k*_cat_ are moderately affected by 3-to 5-fold compared with H4-5.

**Figure 2.**
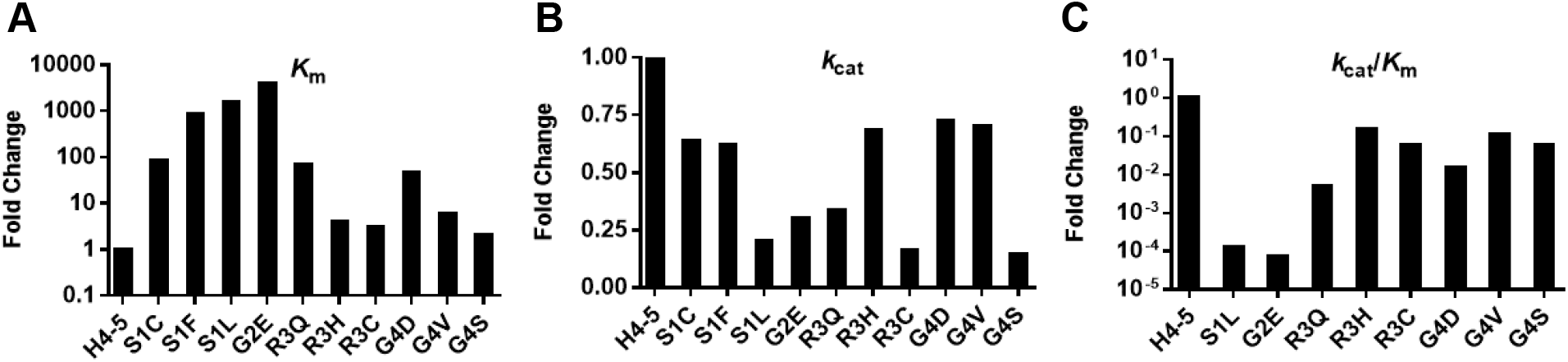
Nα-acetylation of oncohistone H4/H2A peptides by hNatD. Fold change of (A) *K*_m_ (B) *k*_cat_, and (C) *k*_cat_/*K*_m_ normalized to wild-type H4/H2A-5.

Arg-3 contributes to hNatD substrate-specific binding through several interactions with the side chains of Asp-127, Glu-129, Tyr-136, and Tyr-138 of hNatD.^10^ R3H mutant of H2A has been observed in uterine and ovarian carcinomas.^22^ In comparison to H4-5, the *k*_cat_ values were decreased by 3-fold to 3.7±0.12 min^-1^ for R3Q and 1.5-fold to 7.5±0.13 min^-1^ for R3H. Interestingly, the R3H mutant displayed a moderate change in catalytic efficiency by 6.5-fold (*k*_cat_/*K*_m_ = 2.9 × 10^5^ M^-1^ min^-1^). Conversely, a 200-fold reduction in *k*_cat_/*K*_m_ (9.4 × 10^3^ M^-1^ min^-1^) was observed in the R3Q mutant. Similarly, hNatD displayed a 17-fold reduction in catalytic efficiency for the R3C mutant (*k*_cat_/*K*_m_ = 1.1 × 10^5^ M^-1^ min^-1^). This difference in catalytic efficiency between R3Q, R3H, and R3C is consistent with the hNatD mutagenesis study that demonstrated the importance of the positive charge at the third residue for substrate binding.^10^

Like Gly-2, Gly-4 resides in a narrow groove that is tailored for glycine. Mutation of Gly-4 to aspartic acid (G4D) or valine (G4V) resulted in slightly decreased catalytic activities by less than 2-fold (G4D *k*_cat_ = 8.0 ± 0.66 and G4V *k*_cat_ = 7.7 ± 0.13 min^−1^, respectively). Despite the similar catalytic activity, hNatD displayed a 7-fold higher catalytic efficiency for G4V (2.1 × 10^4^ M-^1^ min^-1^) than G4D (2.9 × 10^4^ M^-1^ min^-1^). When Gly-4 was mutated to serine (G4S), hNatD exhibited a 17- and 7-fold reduction in *k*_cat_/*K*_m_ and *k*_cat_ values, respectively. The co-crystal structure of hNatD in complex with its substrate peptide (PDB: 4U9W) indicated that Gly-4 interacts with Trp-90, Thr-174, and Tyr-211.^10^ However, it is unclear why G4S resulted in a salient reduction in *k*_cat_/*K*_m_ and *k*_cat_ values among three mutations.

### Crosstalk between other PTMs and Nα Acetylation

Several PTMs have been demonstrated to be affected by Nα-acetylation at the H4 to induce different gene expression patterns.^25,27,29^ Nα-acH4 in lung cancer activate Slug gene expression through the reducing H4pS1 and enhancing H4R3me2a and H4K5ac.^27^ On the other hand, Nα-acH4 transcriptionally induces PRMT5 expression and thus increases the H4R3me2s in colorectal cancer cells.^29^ To determine how chemical modifications with charged and steric properties on the substrate peptides affect hNatD activity, we synthesized and determined the steady-state kinetics of several synthetic H4 peptides with common modifications (Table 3). Upon phosphorylation of H4S1, hNatD showed a 2.3-fold drop in *k*_cat_ (4.6 ± 0.090 min^−1^) and a 50-fold increase in *K*_m_ (303 ± 21 μM), leading to an over 100-fold reduction in catalytic efficiency. As hNatD has a restrictive size pocket that favors a small side chain, phosphorylation on Ser-1 may not be tolerable in the pocket. In addition, the negatively charged phosphate group may also clash with the Glu-139 of hNatD.^10^ Thus, it is not surprising that phosphorylation at Ser-1 inhibits Nα-acetylation.

**Table 3.**
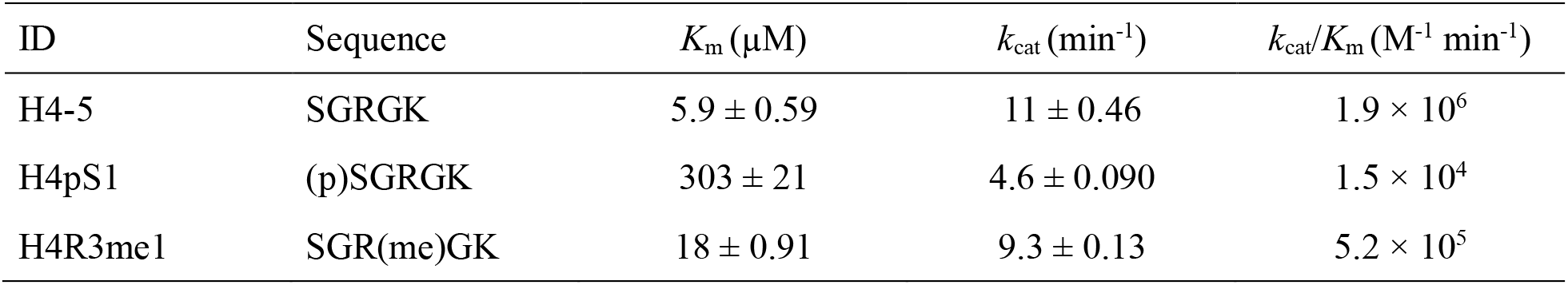
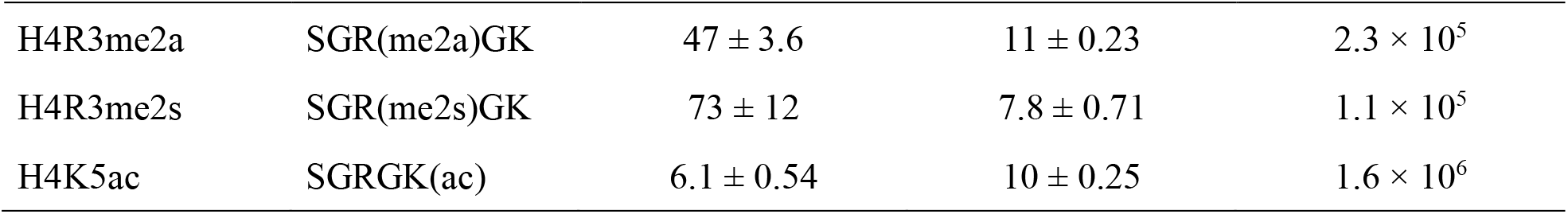
Impact of histone H4S1, H4R3, and H4K5 modification on Nα-acetylation.

The guanidino group of Arg-3 side chain interacts with Asp-127, Glu-129, and Tyr-138 of NatD through hydrogen bonds.^10^ Thus, we set to investigate the effects of Arg-3 methylation on Nα-acetylation. Our biochemical studies showed that hNatD had a similar catalytic activity for methylation of H4R3 despite different methylation states (me1, me2a, or me2s) compared to unmodified H4-5 (Table 3, Figure 3). However, the impacts of methylation states on catalytic efficiency are different. Specifically, hNatD displayed 3.6-fold decreased catalytic efficiency for H4R3me1 with a *k*_cat_/*K*_m_ value of 5.2 × 10^5^ M^-1^ min^-1^, while an 8- and 17-fold reduction were observed for H4R3me2a and H4R3me2s, respectively. As previous biochemical studies reported that Nα-acetylation of H4 slightly reduced PRMT1, 3, 5, and 8 activity on Arg-3 dimethylation, our findings suggest the effects between Nα-acetylation and Arg-3 dimethylation are mutually inhibitory.^28,30^ On the occasion of Lys-5 acetylation (H4K5ac), hNatD exhibited comparable kinetic parameters for H4K5ac as H4-5. One possible explanation is that the side chain of Lys-5 has less contribution to NatD substrate binding because of its exposure to solvent.^10^

**Figure 3.**
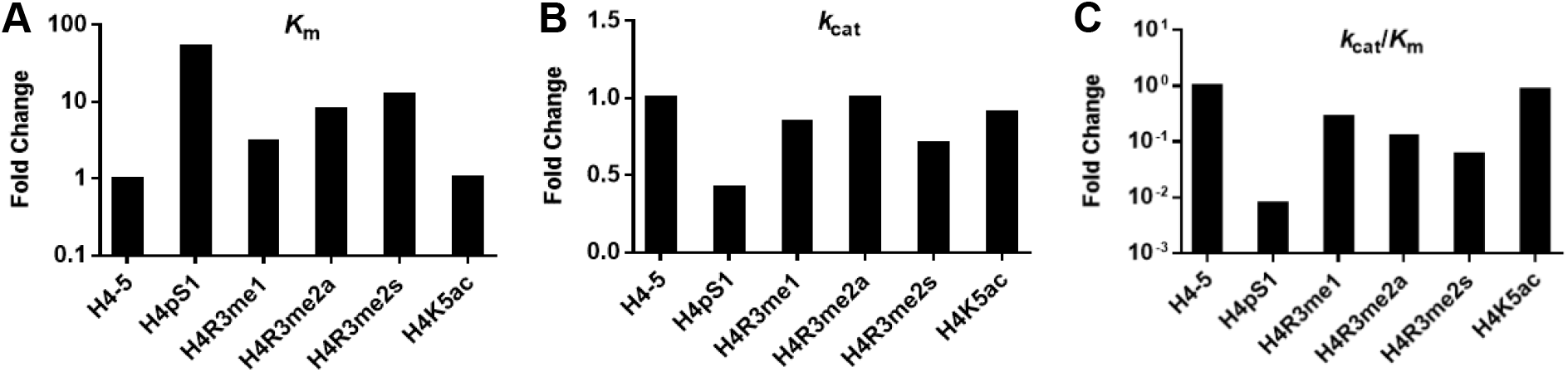
Impact of chemically modified H4 pentapeptides on hNatD-mediated acetylation. Fold change of (A) *K*_m_ (B) *k*_cat_, and (C) *k*_cat_/*K*_m_ normalized to wild-type H4-5.

## Conclusion

In this study, we examined how the length, oncohistone mutations, and PTMs at the substrate recognition motif affect Nα acetylation. Based on our observation, the oncohistone mutations at Ser-1 and Gly-2 of H4 and H2A strongly inhibit hNatD catalysis and hampers substrate recognition by over 100-fold (Figure 4). Compared with mutations at Ser-1 and Gly-2, hNatD was moderately inhibited by most mutations at Arg-3 and Gly-4 but slightly affected by the R3H mutant, which supports the importance of the positive charge status at Arg-3. Meanwhile, our studies provide the first evidence of the negative impact of phosphorylation of Ser-1 and methylation of Arg-3 on Nα-acetylation, uncovering the crosstalk between Nα-acetylation and other PTMs. Specifically, phosphorylation of H4S1 is over 100-fold inhibitory to Nα-acetylation. hNatD displayed a 10–100 fold decreased activity for dimethylation of H4R3 and less than 10-fold reduction in monomethylation of H4R3. The proposed model establishes a relationship between Nα-acetylation and local PTMs of histones and oncohistones, which helps to elucidate the complex crosstalk in histones (Figure 4). This work has provided the mechanistic understanding of how single mutation of oncohistone and adjacent PTMs affect Nα-acetylation by NatD, outlining the basis for future investigation on the functions of the NatD-mediated pathway.

**Figure 4.**
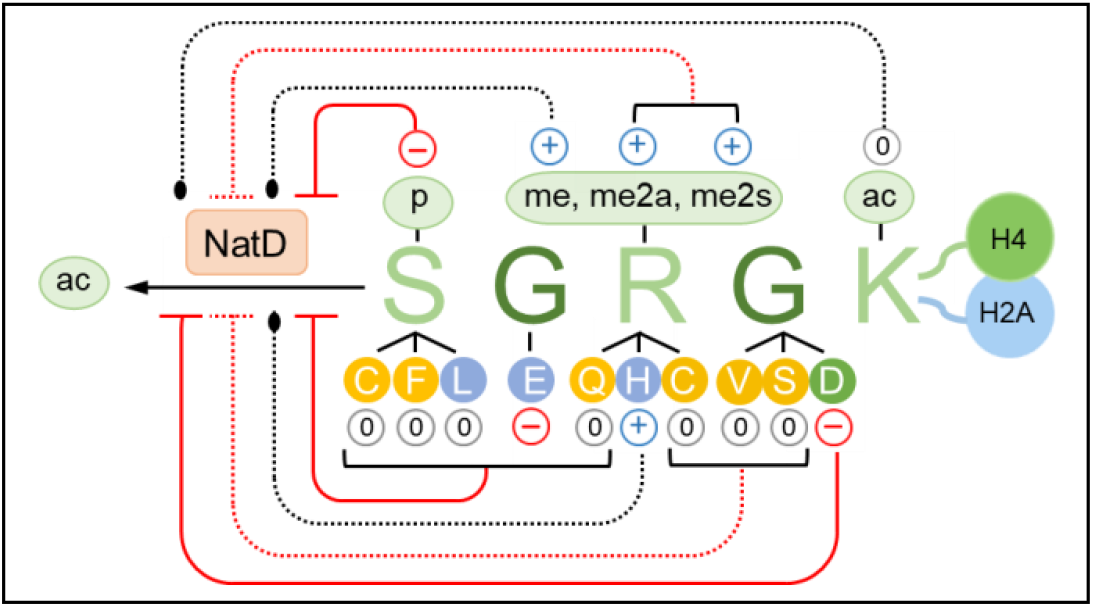
Model for summarizing the impact of oncohistone H4/H2A and PTMs of histone H4 on NatD activity. PTMs include phosphorylation (p), monomethylation (me), asymmetric dimethylation (me2a), symmetric dimethylation (me2s), and acetylation (ac). Mutations were detected on H4 and H2A (yellow), only on H4 (green), only on H2A (blue). The charge status contains neutral (0 within in a gray circle), positive (+ within in a blue circle), and negative (- within in a red circle). Solid red line with blunt end = over 100-fold inhibition, red dotted line =10– 100-fold inhibition, and black dotted line = less than 10-fold inhibition.

### Experimental section

#### Chemical Reagents

The standard Fmoc-protected amino acids were purchased from CEM Corporation. Acetyl coenzyme A lithium salt was purchased from Sigma-Aldrich. High-performance liquid chromatography (HPLC) grade solvents were purchased from Fisher Scientific. Matrix-assisted laser desorption/ionization (MALDI) spectra were acquired using positive-ion mode on a 4800 MALDI TOF/TOF mass spectrometry (Sciex). Peptides were synthesized on a CEM Liberty Blue peptide synthesizer. The purity of the peptides was confirmed by Agilent 1260 Series HPLC system, eluting with a 0–40% acetonitrile/water gradient with 0.1% TFA. All the purity of final peptides showed >95%. The spectra of MALDI and HPLC are available in the Supporting information (Figure S3).

#### Peptide synthesis

The peptides were synthesized using Rink Amide MBHA resin (0.05 mmol) and followed a standard Fmoc protocol. Fmoc protected amino acids were coupled at 0.2 M in DMF using 0.5 M of DIC and 1.0 M of Oxyma. Double coupling of Fmoc-Ser(HPO_3_Bzl)-OH (ChemPep), Fmoc-Arg(me)(Pbf)-OH (AnaSpec), Fmoc-Arg(me)_2_-OH (asymmetrical and symmetrical) (AnaSpec), Fmoc-Lys(ac)-OH (Creosalus) were performed. Fmoc deprotection was carried out by 20% piperidine in DMF. The resin was washed with CH_2_Cl_2_ (5 mL) and MeOH (5 mL) alternatively three times, and the peptide was cleaved by a cleavage cocktail (TFA/TIPS/water/DODT = 94/1/2.5/2.5 v/v) (5 mL) for 0.05 mmol scale of resin for 4 h. The peptide suspension was filtered, and the filtrate was dried under N_2_ and precipitated with cold ether (10 mL). The peptide precipitation was pelleted by centrifugation at 4,200 rpm for 10 minutes, and the resulting supernatant was discarded. The crude peptide pellet was dissolved in deionized water (10 mL), passed through a 0.45 μm filter, and purified by preparative reversed-phase high-performance liquid chromatography (RP-HPLC) on an Agilent 1260 Series system with a C18 column (5 μm, 10 mm × 250 mm) at a flow rate of 4.0 mL/min. Peptides were purified by two mobile phases consisting of 0.1% TFA in water and acetonitrile in a linear gradient.

#### Protein Expression and Purification

Expression and purification of hNatD were carried out as previously described.^31^ Histone H4 protein was a gift from Dr. Chongli Yuan from Purdue University.

#### Continuous fluorescence-based acetylation assay

A fluorescence assay was adapted to study the kinetics of NatD, which monitors the formation of a ThioGlo4-thiol adduct that exhibits a strong fluorescence at 465 nm.^33,34^ NatD activity was measured at 25 °C under the following conditions in a final reaction volume of 40 μL: 50 nM NatD, 15 μM ThioGlo4, and 50 μM AcCoA in reaction buffer (25 mM HEPES, 150 mM NaCl, 0.01% Triton, pH 7.5). For AcCoA characterization, H4-8 was fixed at 50 μM. All components except for peptide substrate were mixed in a volume of 36 μL and incubated at 25 °C for 10 min. The reaction was initiated by adding 4 μL of varying concentrations of peptide substrates. Fluorescence was monitored on a BMG CLARIOstar microplate reader with excitation of 400–415 nm and 460–485 nm. For quantifying the fluorescent signal, a previously reported standard curve (y = 5176x + 344.6) was applied to convert arbitrary fluorescence units to the concentration of CoA.^31^ GraphPad Prism 7 was used to fit the initial velocity to Michaelis-Menten or Substrate inhibition model. The experiments were performed in at least duplicate.

#### MALDI-MS acetylation assay

Kinetics of CGRGK and SGCGK were characterized with a MALDI-MS assay at 25 °C under the following conditions in a final reaction volume of 40 μL: 50 nM of NatD and 50 μM of AcCoA in reaction buffer (25 mM HEPES, 150 mM NaCl, pH 7.5). The 10 mM cysteine peptides (CGRGK and SGCGK) were allowed to reduce under 50 mM of DTT at room temperature for 30 min. The peptides were then 2-fold diluted from 1 mM (CGRGK) and 400 μM (SGCGK) with dilution buffer (25 mM HEPES, 150 mM NaCl, 10 mM DTT, pH 7.5). All components except for peptide substrate were mixed in a volume of 36 μL, and the reaction was initiated with 4 μL of varying concentrations of peptide substrates. At desired time points, aliquots were quenched in a 1:1 ratio with the 0.1% of TFA aqueous solution and spotted with matrix solution (10 mg/ml of CHCA in water/acetonitrile/TFA = 1:1:0.1% v/v/v) on the MALDI plate. The initial velocity was determined by calculating the slope from aliquots taken at differing time points. GraphPad Prism 7 was used to fit the initial velocity to Michaelis-Menten or Substrate inhibition model. The experiments were performed in duplicate.

## ASSOCIATED CONTENT

### Supporting Information

The following files are available free of charge.

Kinetic profiles, MALDI-MS, and HPLC spectra of all peptides.

## AUTHOR INFORMATION

### Author Contributions

The manuscript was written through contributions of both authors. Both authors have given approval to the final version of the manuscript.

### Funding Sources

We thank the support from the Department of Medicinal Chemistry and Molecular Pharmacology (RH).

### Notes

The authors declare no competing financial interest.

## ACKNOWLEDGMENT

We thank Mr. Yuanrui Zhao for his assistance with several peptide synthesis and Dr. Chongli Yuan for H4 protein.

## ABBREVIATIONS

NATs: N-terminal acetyltransferase
AcCoA: acetyl-coenzyme A
H4pS1: histone H4 serine 1 phosphorylation
H4R3me2a: histone H4 arginine 3 asymmetric dimethylation
H4R3me1: histone arginine 3 monomethylation
H4R3me2a: histone H4 arginine 3 symmetric dimethylation
H4K5ac: histone H4 lysine 5 acetylation;

## Notes

### Competing Interest Statement

The authors have declared no competing interest.

