## Supporting information for "Effects of Oncohistone Mutations and PTM Crosstalk on the N-terminal Acetylation Activities of NatD"

### Table of Contents

Figure S1 Kinetic profiles of peptides by MALDI-MS assay ..................................................S2
Figure S2 Kinetic profiles of peptides by fluorescence assay ..................................................S3
Figure S3 MALDI-MS and HPLC spectra of peptides..............................................................S4

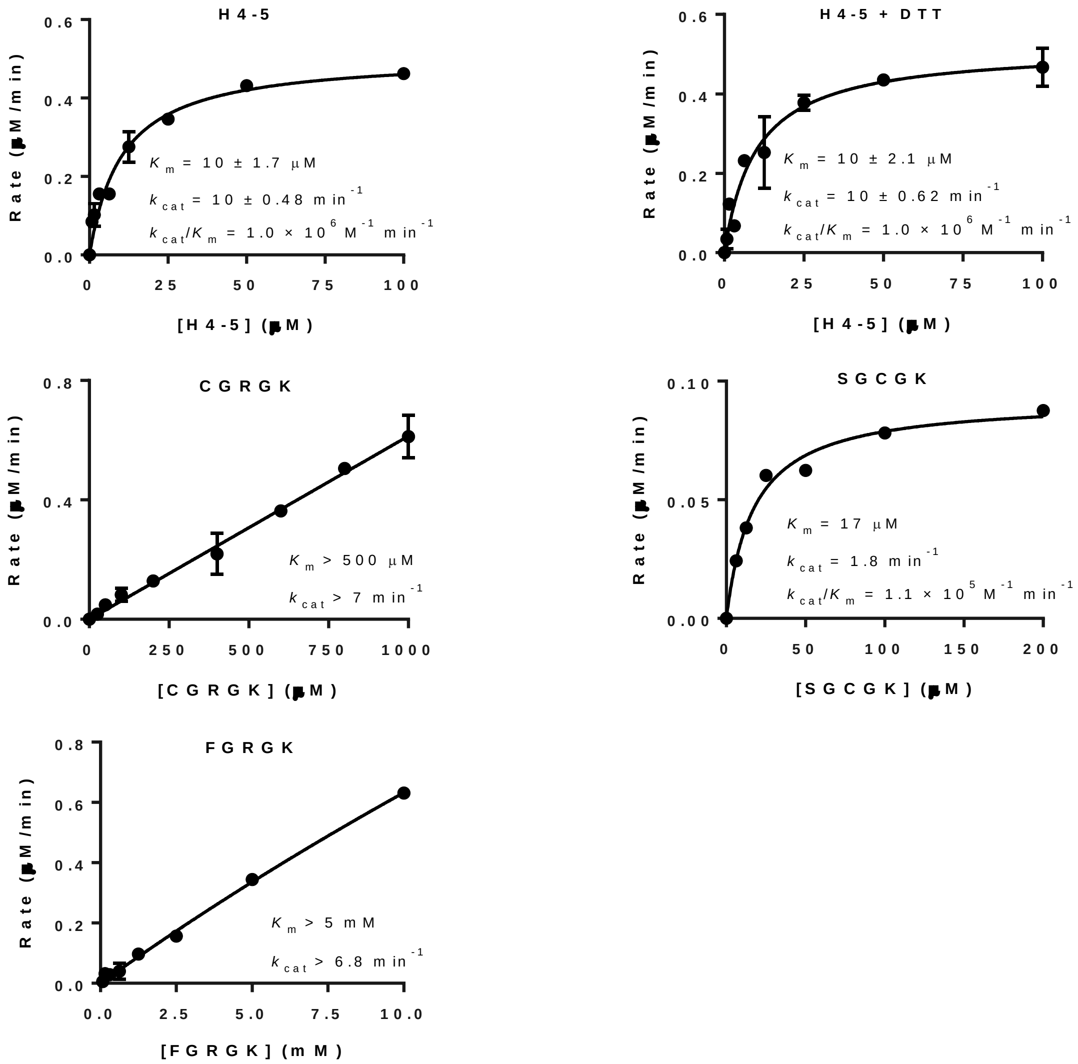


**Figure S1.** Kinetic profiles of peptides of (A) H4-5, (B) H4-5 with 5 mM of DTT, (C) CGRGK with 5 mM DTT, (D) SGCGK with 1 mM of DTT, and (D) FGRGK. Single point of MALDI-MS acetylation assay was performed with 50 nM of hNatD, 50 µM of AcCoA, and varying concentrations of peptide substrate at 25 ºC. All experiments were performed in duplicate except for SGCGK, which was performed in a single trial.

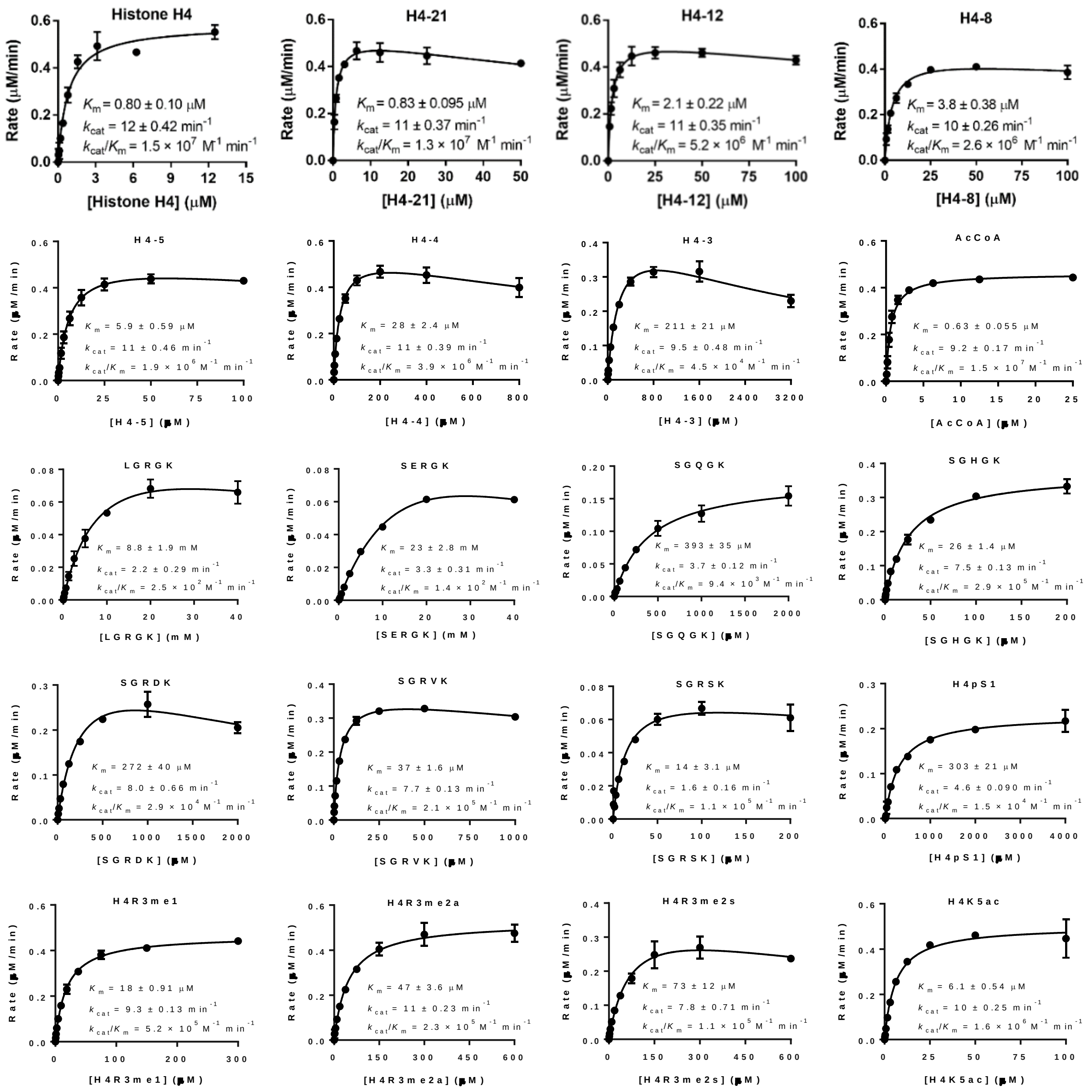


**Figure S2.** Kinetic profiles of (A) histone H4, (B) H4-21, (C) H4-12, (D) H4-8, (E) H4-5, (F) H4-4, (G) H4-3, (H) AcCoA, (I) LGRGK, (J) SERGK, (K) SGQGK, (L) SGHGK, (M) SGRDK, (N) SGRVK, (O) SGRSK, (P) H4pS1, (Q) H4R3me1, (R) H4R3me2a, (S) H4R3me2s, and (T) H4K5ac determined by the fluorescence assay. For peptide kinetics, the assay was performed with 50 nM hNatD, 50 µM of AcCoA, and varying concentrations of peptide substrate at 25 ºC. For AcCoA kinetics, the assay was performed with 50 nM hNatD, 50 µM H4-8, and varying concentrations of AcCoA at 25 ºC. All experiments were performed in at least duplicate.

**Figure S3.** MALDI-MS and HPLC spectra of peptides

**H4-3**

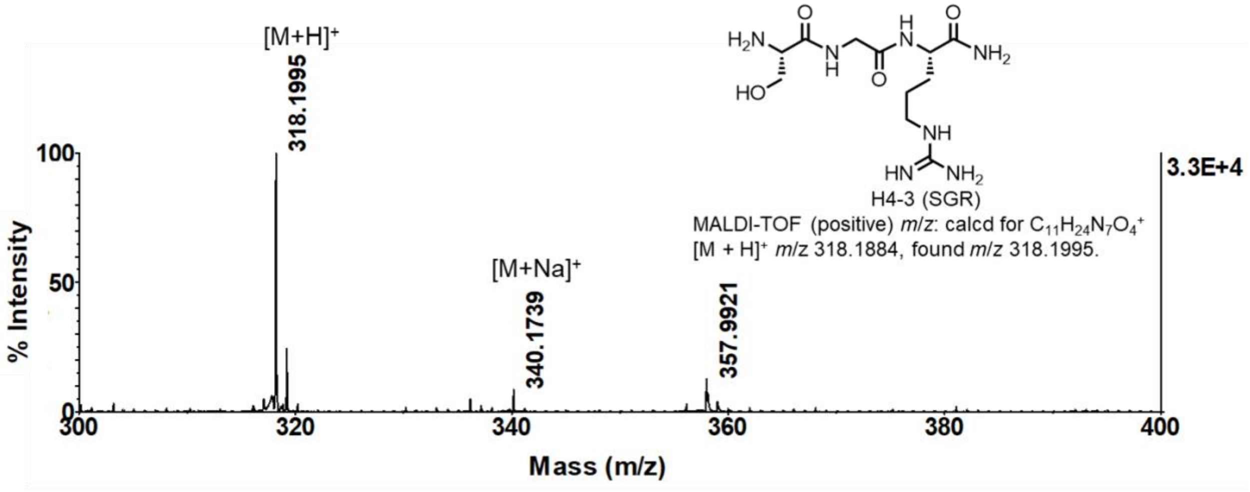


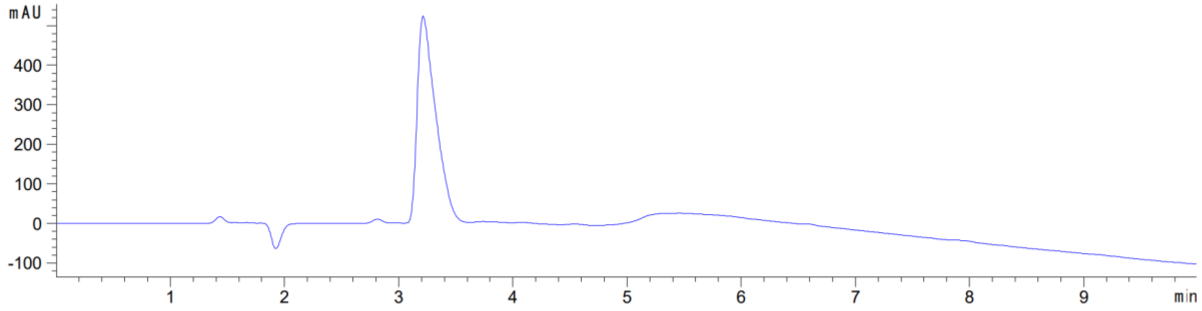


**H4-4**

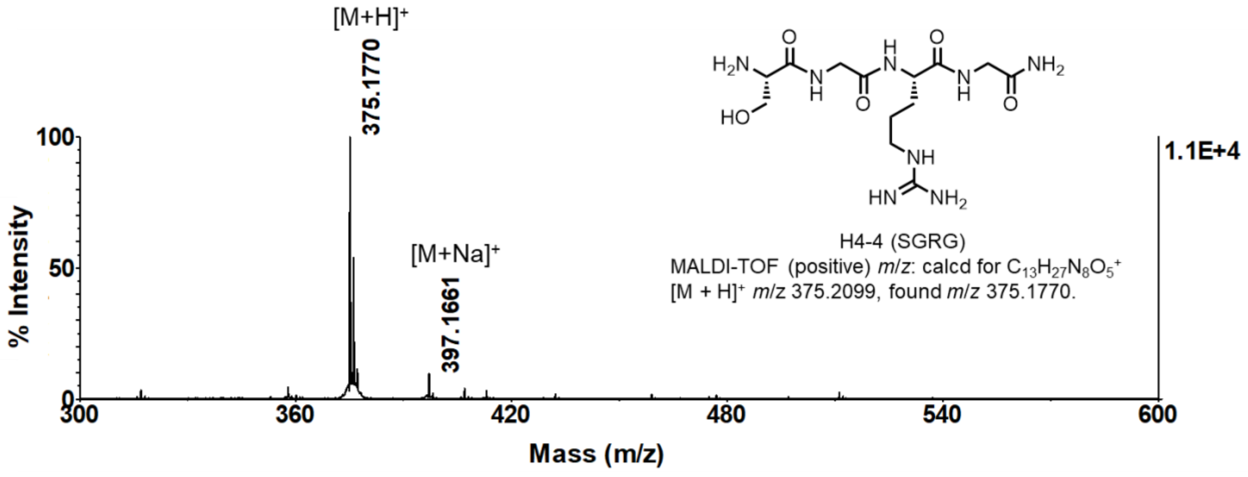


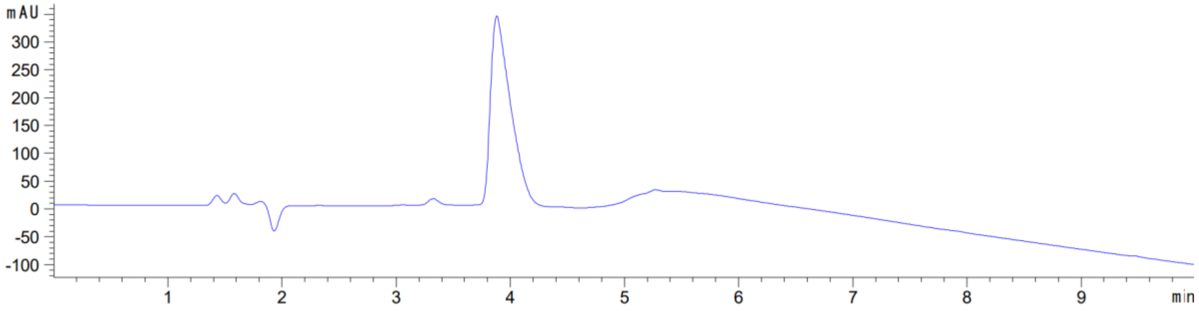

**H4-5**

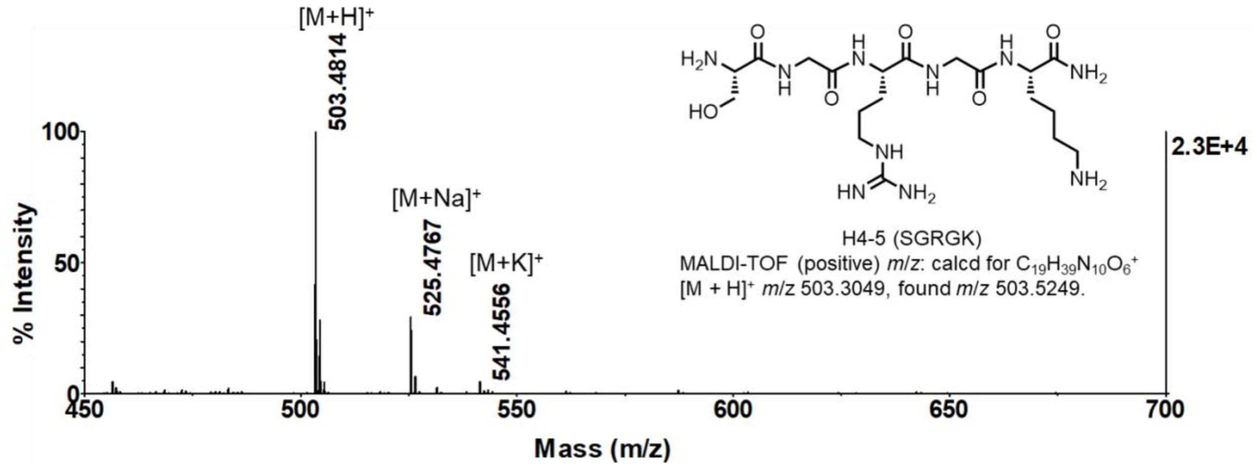
[M+H]⁺
503.4814
[M+Na]⁺
525.4767
[M+K]⁺
541.4556
% Intensity
Mass (m/z)
2.3E+4
H4-5 (SGRGK)
MALDI-TOF (positive) *m/z*: calcd for $C_{19}H_{39}N_{10}O_6^+$
[M + H]⁺ *m/z* 503.3049, found *m/z* 503.5249.

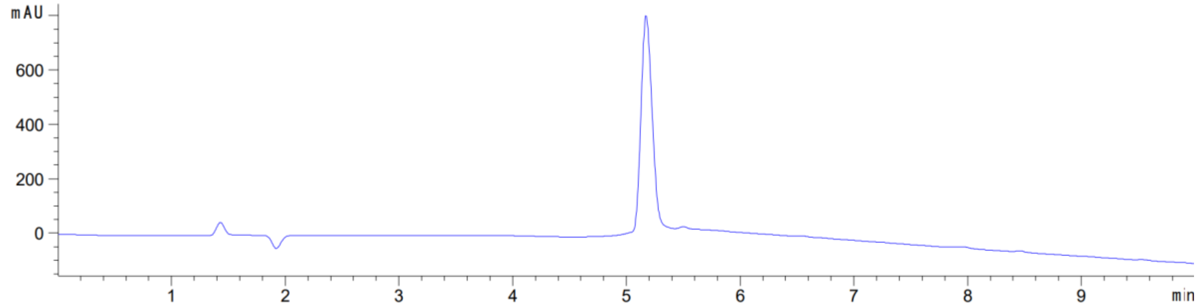
mAU
min

**H4-8**

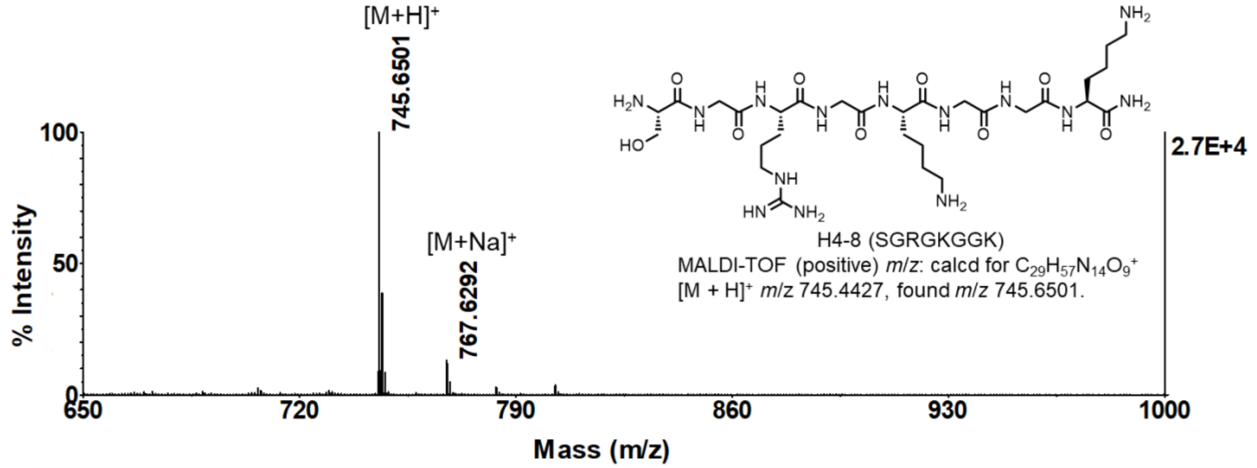
[M+H]⁺
745.6501
[M+Na]⁺
767.6292
% Intensity
Mass (m/z)
2.7E+4
H4-8 (SGRGKGGK)
MALDI-TOF (positive) *m/z*: calcd for $C_{29}H_{57}N_{14}O_9^+$
[M + H]⁺ *m/z* 745.4427, found *m/z* 745.6501.

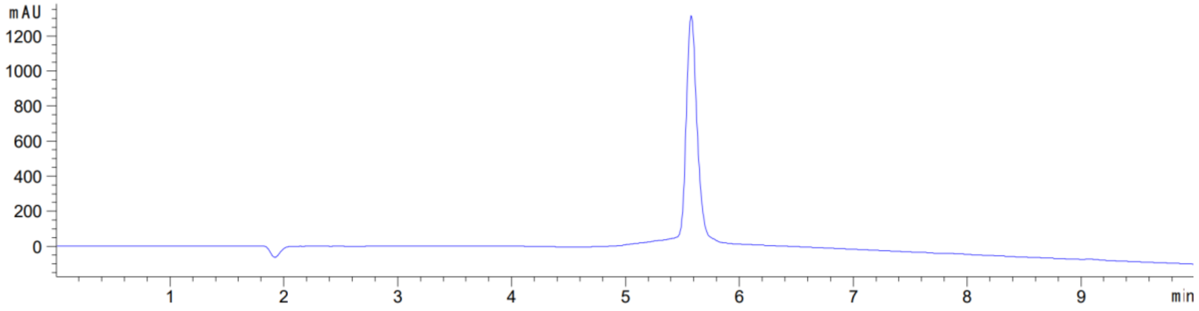
mAU
min

**H4-12**

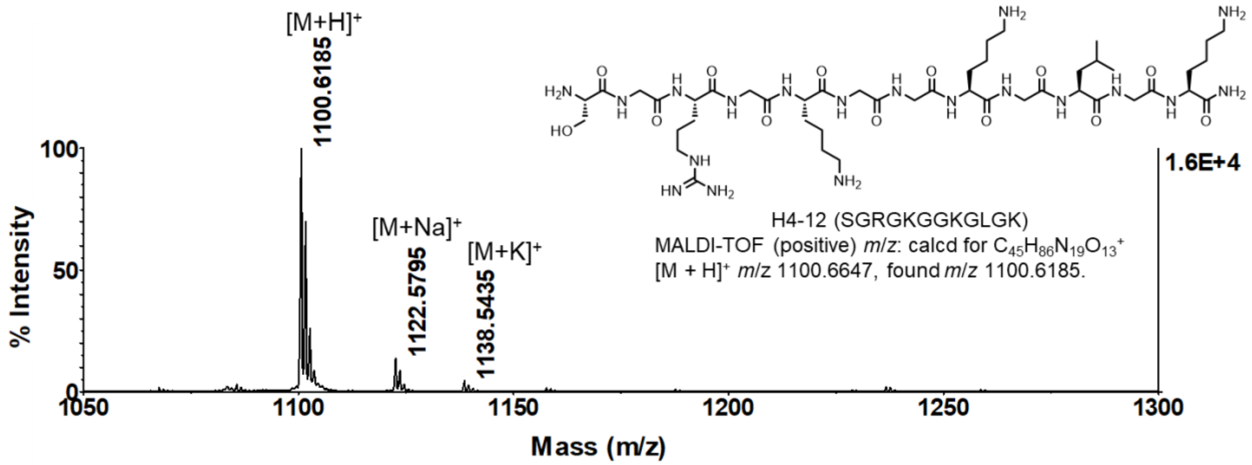
[M+H]+
1100.6185
[M+Na]+
1122.5795
[M+K]+
1138.5435
1.6E+4
H4-12 (SGRGKGGKGLGK)
MALDI-TOF (positive) m/z: calcd for $C_{45}H_{86}N_{19}O_{13}^+$
[M + H]+ m/z 1100.6647, found m/z 1100.6185.
% Intensity
Mass (m/z)

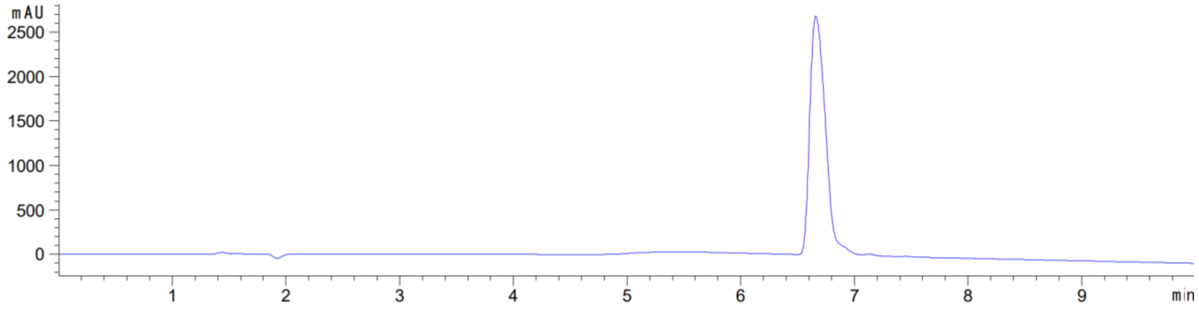
mAU
min

**H4-21**

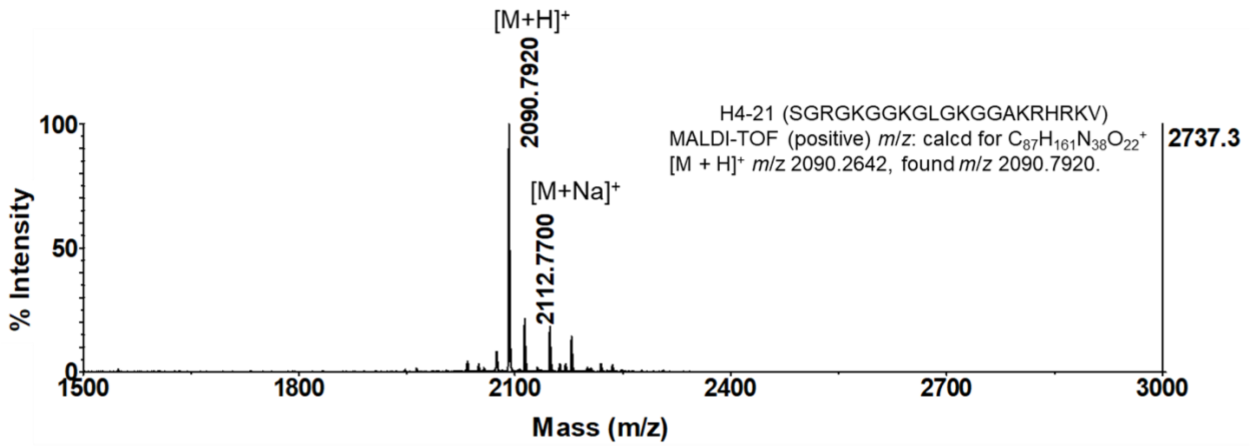
[M+H]+
2090.7920
[M+Na]+
2112.7700
H4-21 (SGRGKGGKGLGKGGAKRHRKV)
MALDI-TOF (positive) m/z: calcd for $C_{87}H_{161}N_{38}O_{22}^+$
[M + H]+ m/z 2090.2642, found m/z 2090.7920.
2737.3
% Intensity
Mass (m/z)

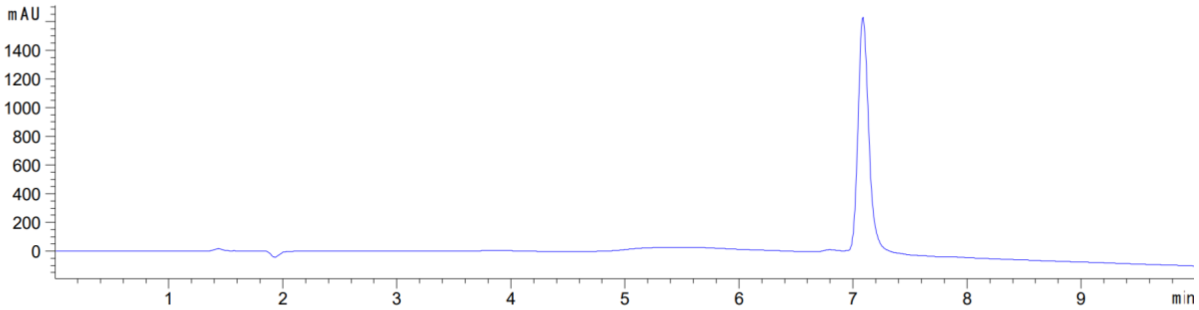
mAU
min

**CGRGK**

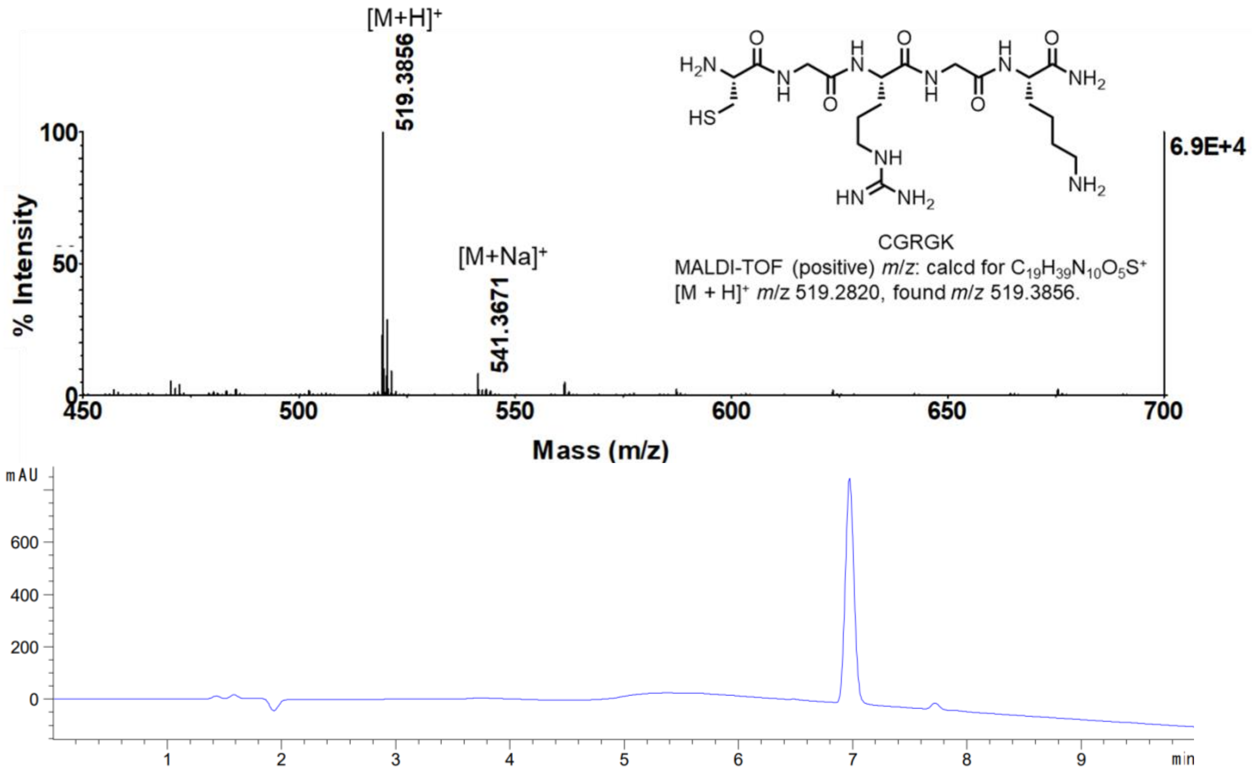

[M+H]+
519.3856
[M+Na]+
541.3671
100
50
0
% Intensity
6.9E+4
450
500
550
600
650
700
Mass (m/z)
CGRGK
MALDI-TOF (positive) m/z: calcd for C19H39N10O5S+
[M + H]+ m/z 519.2820, found m/z 519.3856.
mAU
600
400
200
0
1
2
3
4
5
6
7
8
9
min


**LGRGK**

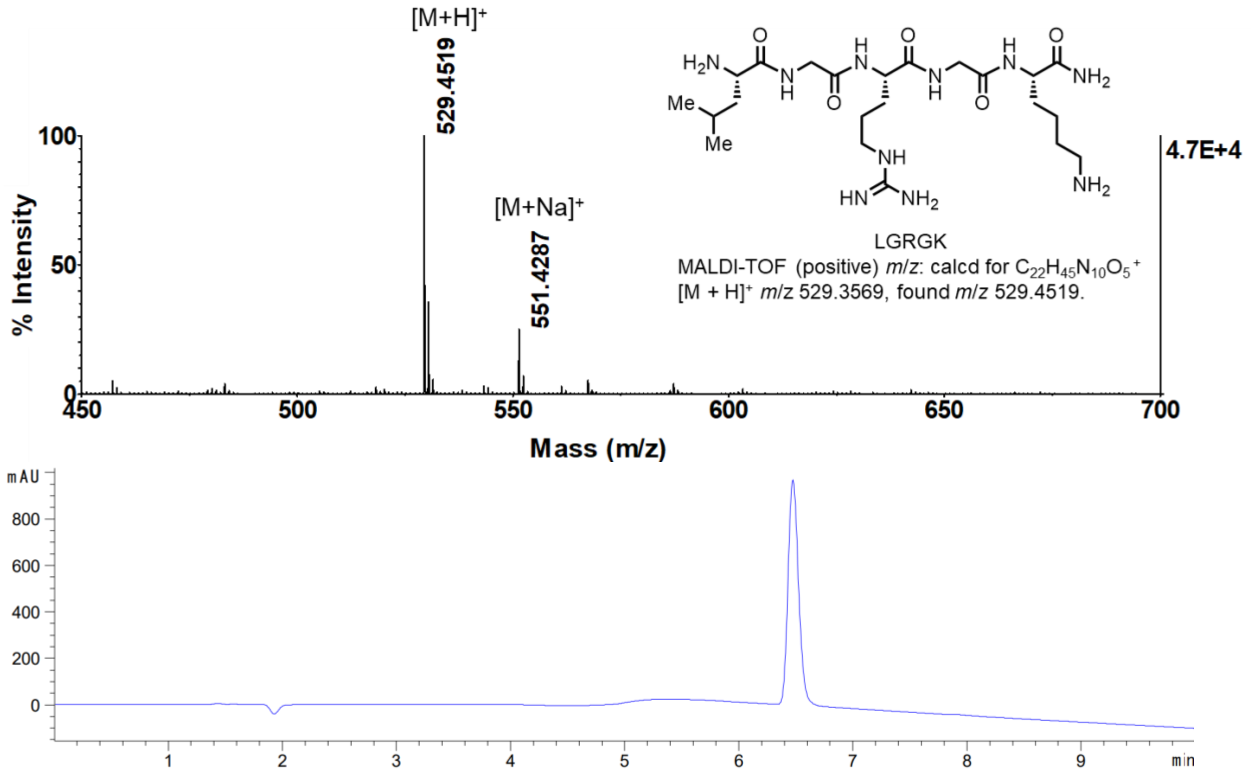

[M+H]+
529.4519
[M+Na]+
551.4287
100
50
0
% Intensity
4.7E+4
450
500
550
600
650
700
Mass (m/z)
LGRGK
MALDI-TOF (positive) m/z: calcd for C22H45N10O5+
[M + H]+ m/z 529.3569, found m/z 529.4519.
mAU
800
600
400
200
0
1
2
3
4
5
6
7
8
9
min

**FGRGK**

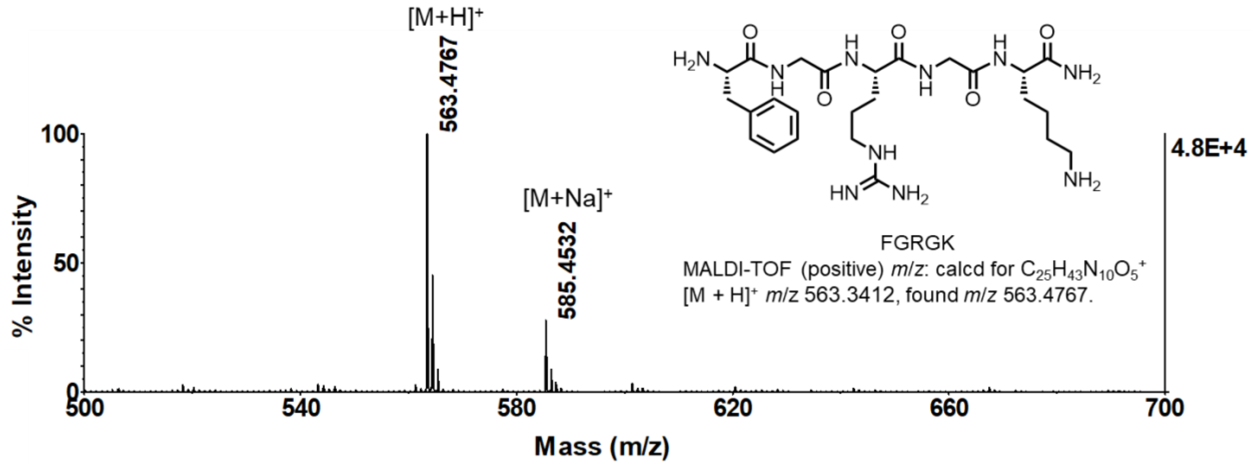

[M+H]+
563.4767
[M+Na]+
585.4532
100
50
0
% Intensity
500
540
580
620
660
700
Mass (m/z)
4.8E+4
FGRGK
MALDI-TOF (positive) *m/z*: calcd for $C_{25}H_{43}N_{10}O_5^+$
[M + H]+ *m/z* 563.3412, found *m/z* 563.4767.


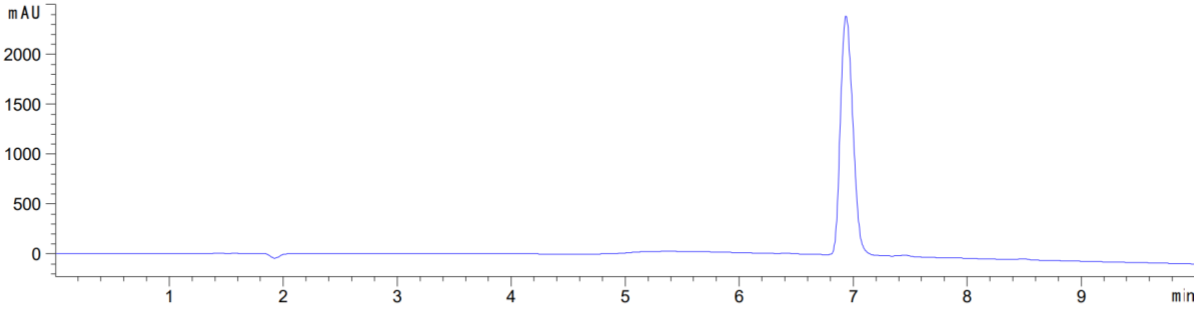

mAU
2000
1500
1000
500
0
1
2
3
4
5
6
7
8
9
min


**SERGK**

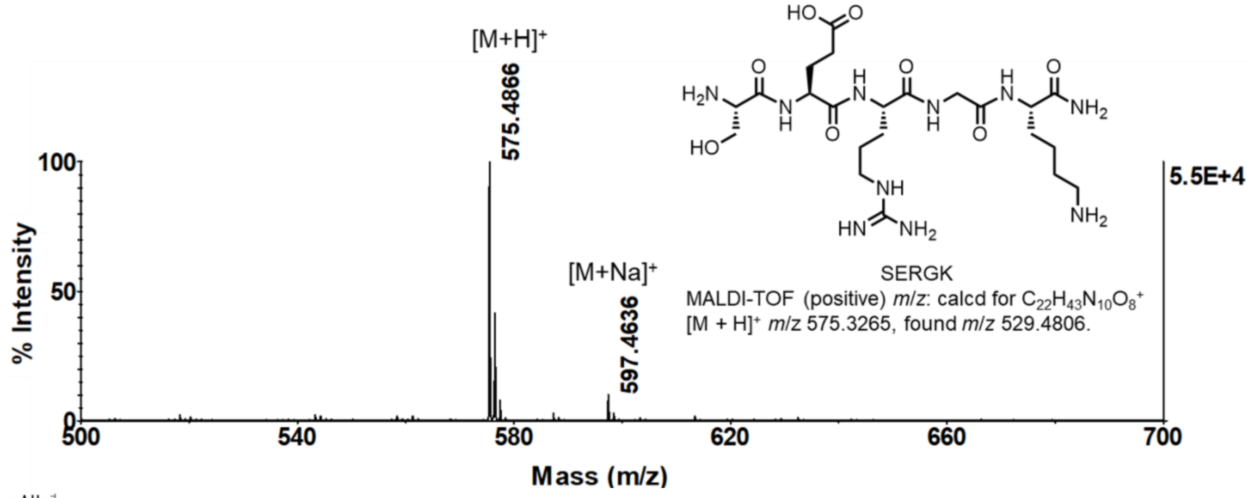

[M+H]+
575.4866
[M+Na]+
597.4636
100
50
0
% Intensity
500
540
580
620
660
700
Mass (m/z)
5.5E+4
SERGK
MALDI-TOF (positive) *m/z*: calcd for $C_{22}H_{43}N_{10}O_8^+$
[M + H]+ *m/z* 575.3265, found *m/z* 529.4806.


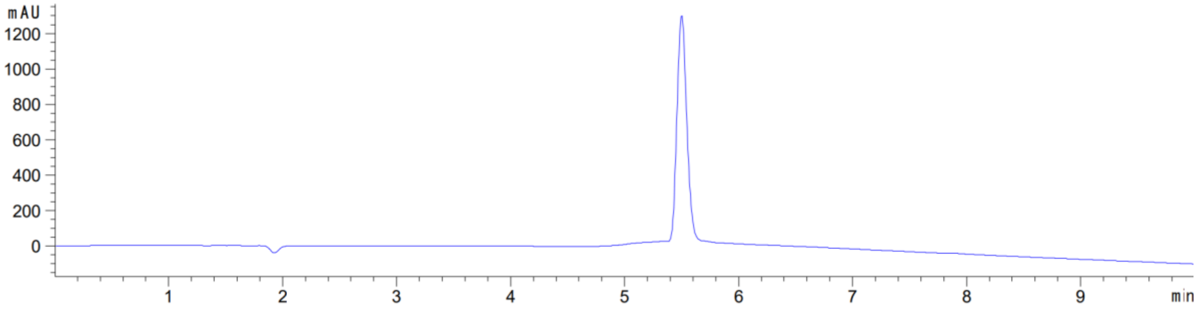

mAU
1200
1000
800
600
400
200
0
1
2
3
4
5
6
7
8
9
min

**SGCGK**

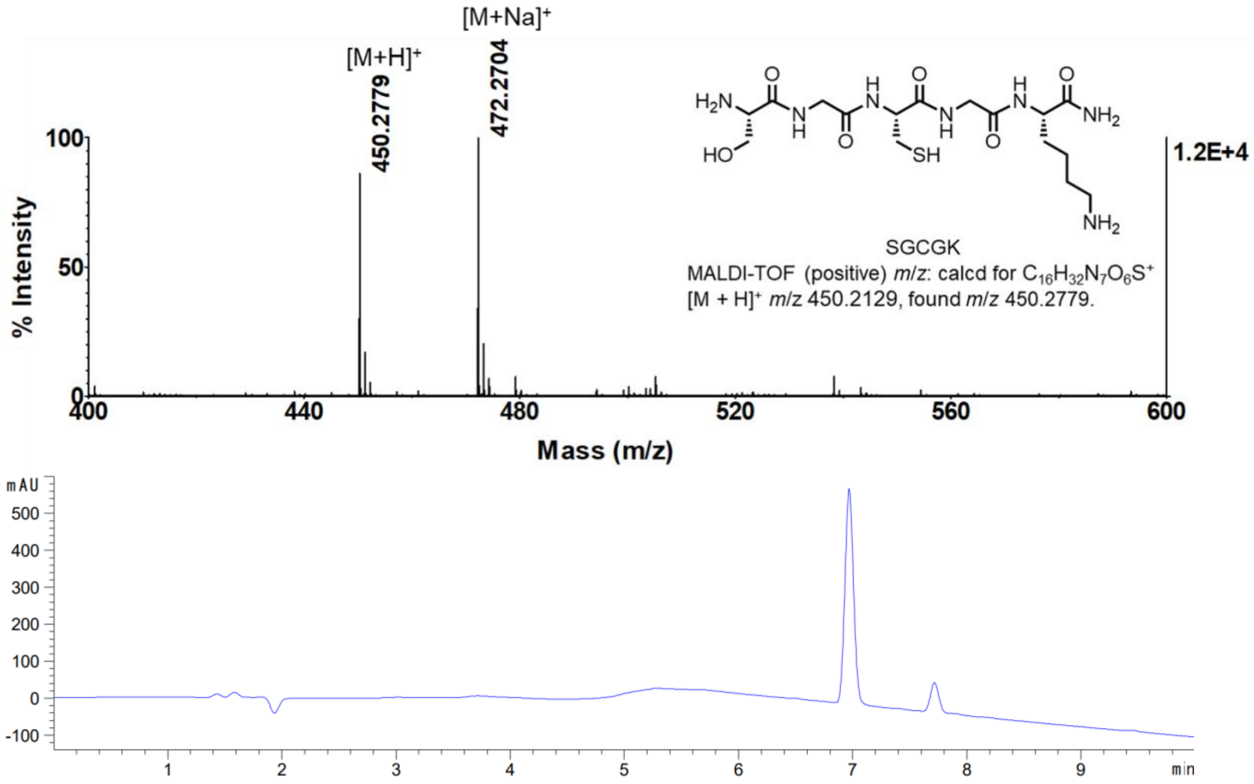
[M+H]+
450.2779
[M+Na]+
472.2704
% Intensity
Mass (m/z)
1.2E+4
SGCGK
MALDI-TOF (positive) m/z: calcd for $C_{16}H_{32}N_7O_6S^+$
[M + H]+ m/z 450.2129, found m/z 450.2779.
mAU
min

**SGHGK**

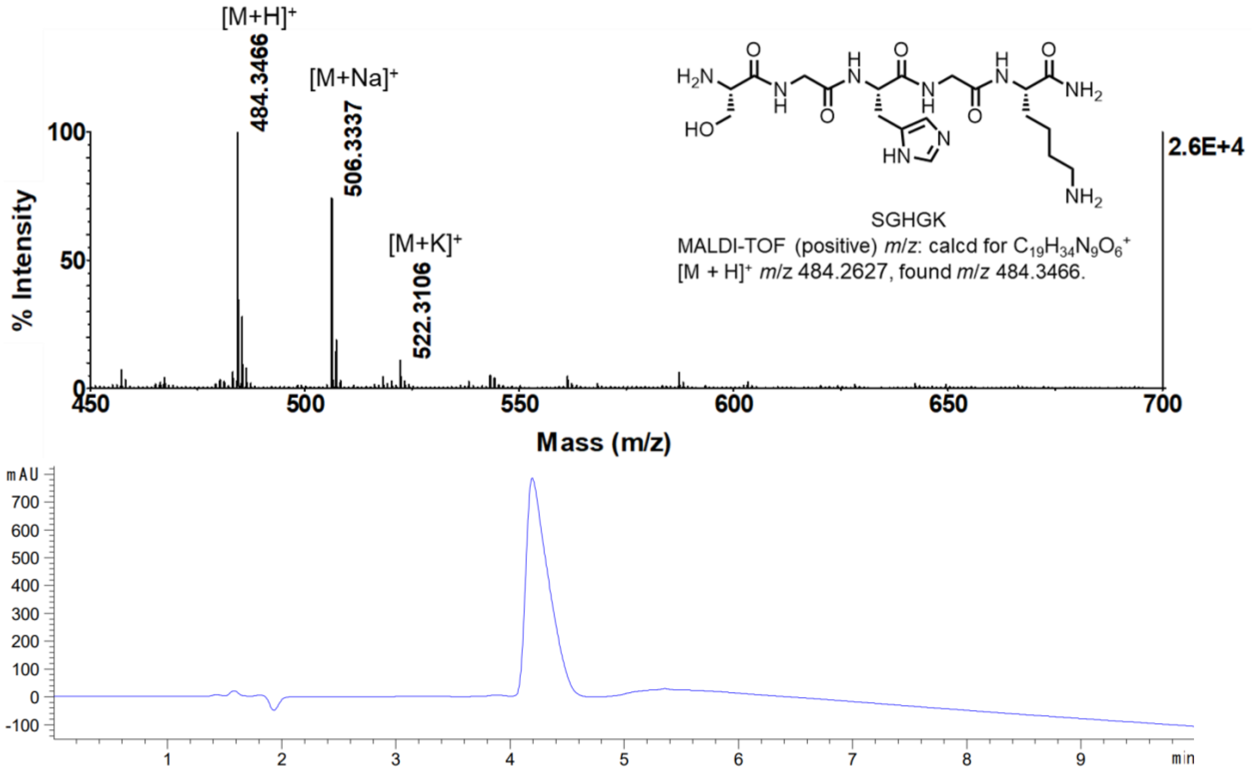
[M+H]+
484.3466
[M+Na]+
506.3337
[M+K]+
522.3106
% Intensity
Mass (m/z)
2.6E+4
SGHGK
MALDI-TOF (positive) m/z: calcd for $C_{19}H_{34}N_9O_6^+$
[M + H]+ m/z 484.2627, found m/z 484.3466.
mAU
min

**SGQGK**

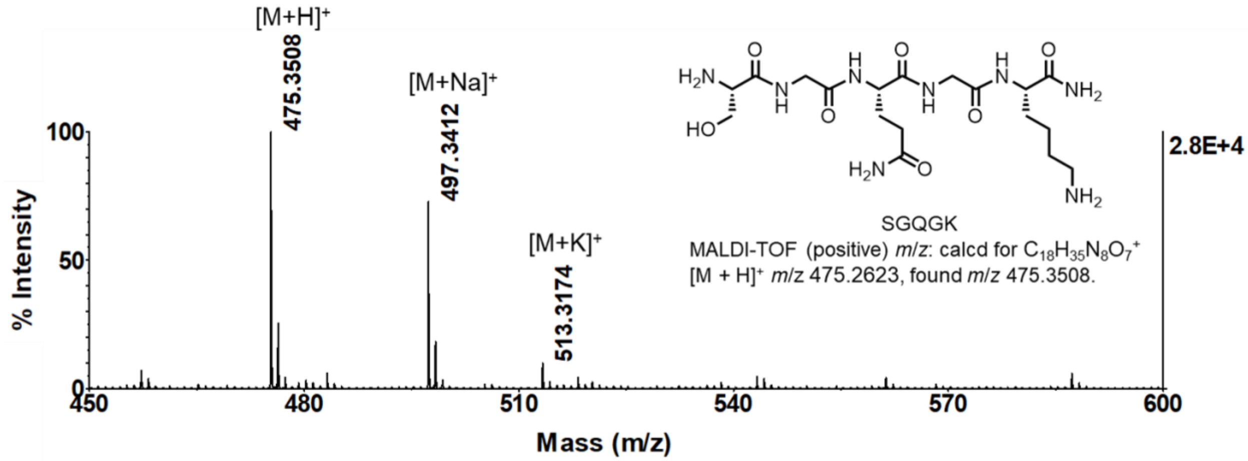

[M+H]+
475.3508
[M+Na]+
497.3412
[M+K]+
513.3174
100
50
0
% Intensity
2.8E+4
450
480
510
540
570
600
Mass (m/z)
SGQGK
MALDI-TOF (positive) m/z: calcd for $C_{18}H_{35}N_8O_7^+$
[M + H]+ m/z 475.2623, found m/z 475.3508.


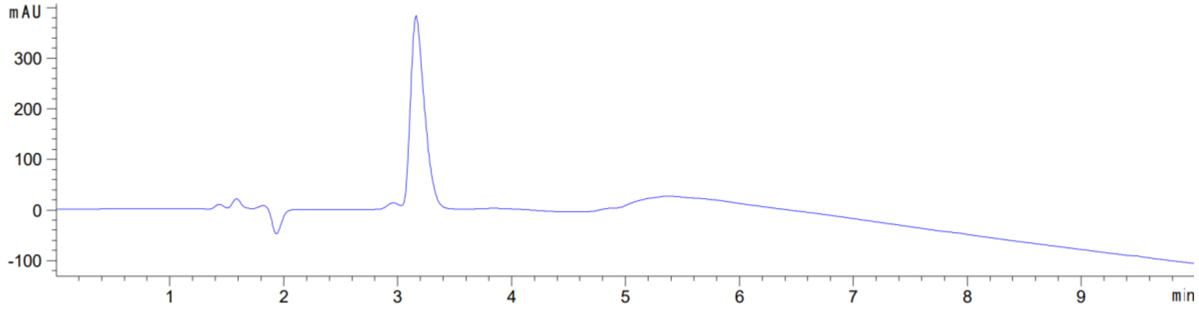

mAU
300
200
100
0
-100
1
2
3
4
5
6
7
8
9
min


**SGRVK**

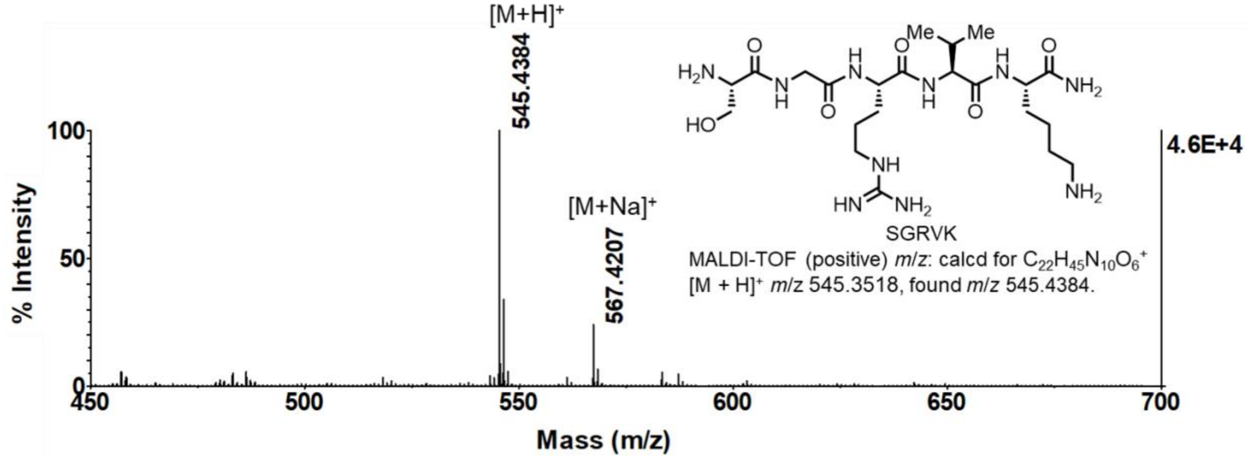

[M+H]+
545.4384
[M+Na]+
567.4207
100
50
0
% Intensity
4.6E+4
450
500
550
600
650
700
Mass (m/z)
SGRVK
MALDI-TOF (positive) m/z: calcd for $C_{22}H_{45}N_{10}O_6^+$
[M + H]+ m/z 545.3518, found m/z 545.4384.


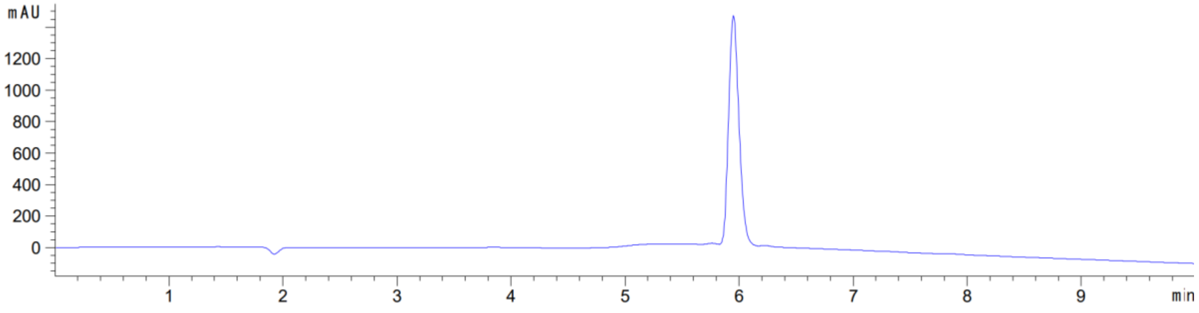

mAU
1200
1000
800
600
400
200
0
1
2
3
4
5
6
7
8
9
min

**SGRSK**

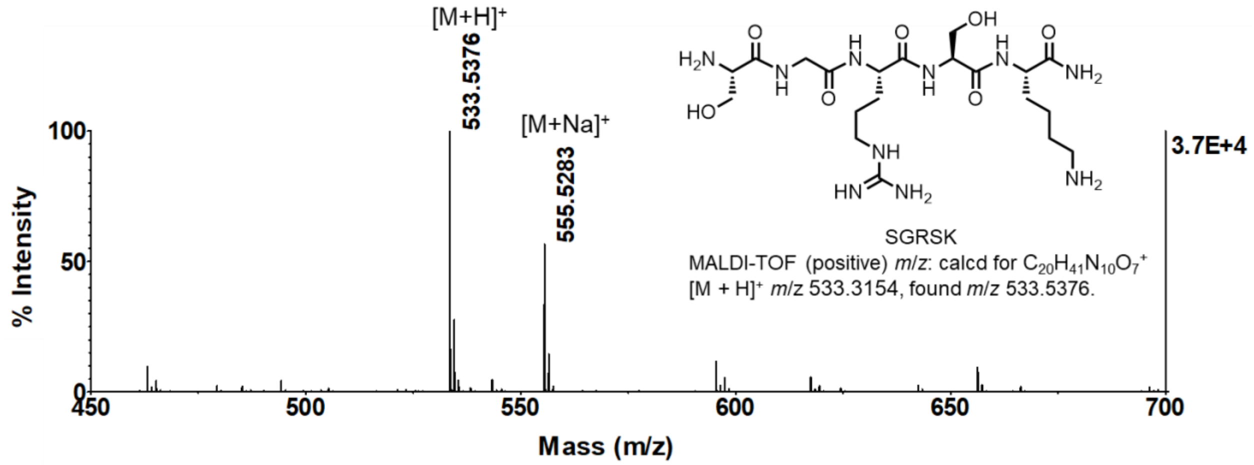
[M+H]+
533.5376
[M+Na]+
555.5283
SGRSK
MALDI-TOF (positive) m/z: calcd for C20H41N10O7+
[M + H]+ m/z 533.3154, found m/z 533.5376.
3.7E+4
% Intensity
Mass (m/z)

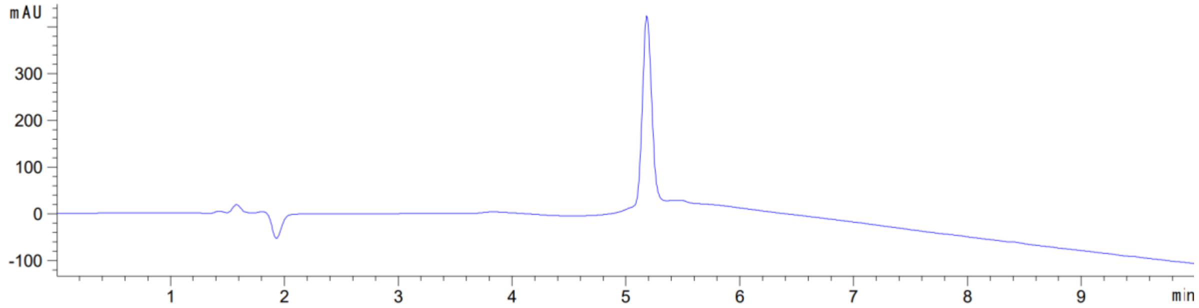
mAU
min

**SGRDK**

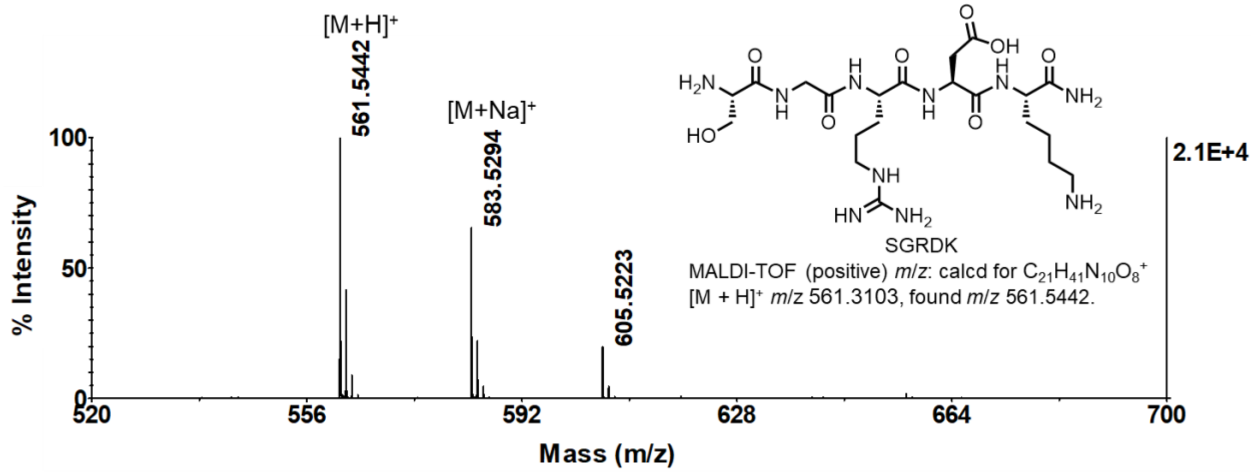
[M+H]+
561.5442
[M+Na]+
583.5294
605.5223
SGRDK
MALDI-TOF (positive) m/z: calcd for C21H41N10O8+
[M + H]+ m/z 561.3103, found m/z 561.5442.
2.1E+4
% Intensity
Mass (m/z)

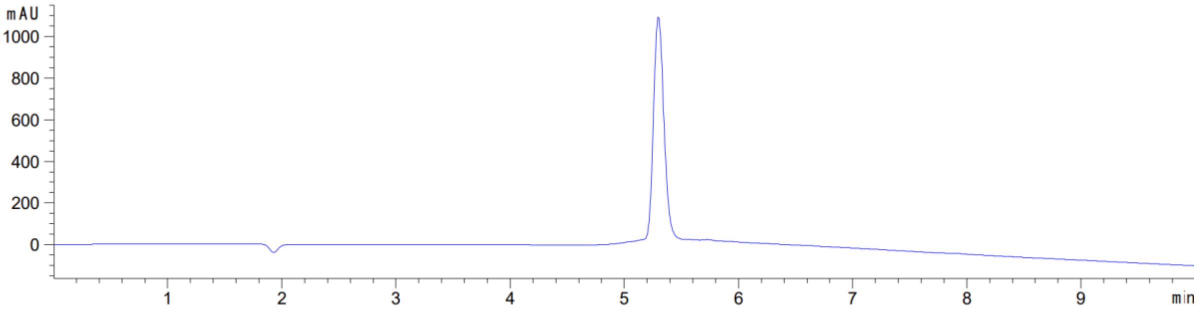
mAU
min

**H4pS1**

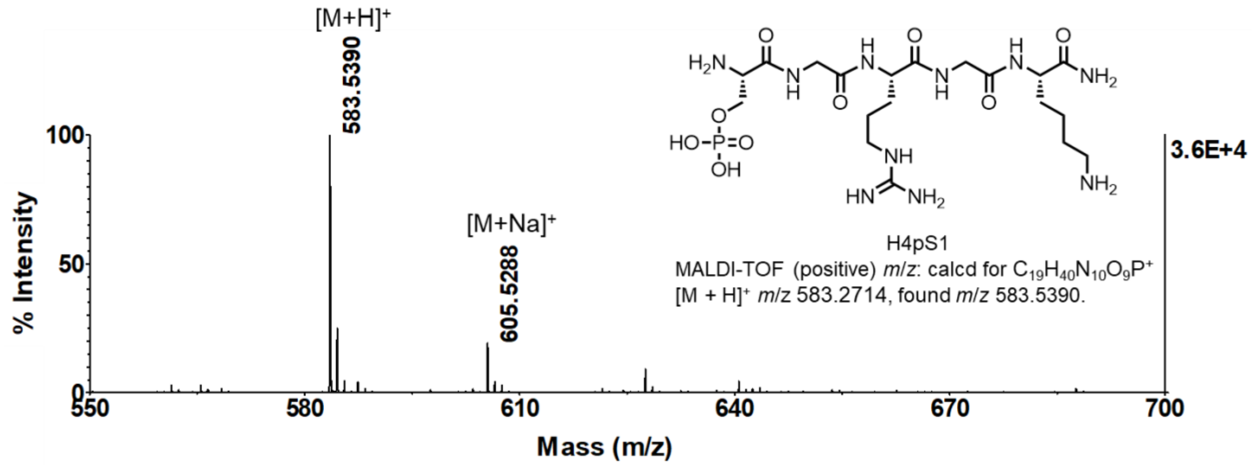


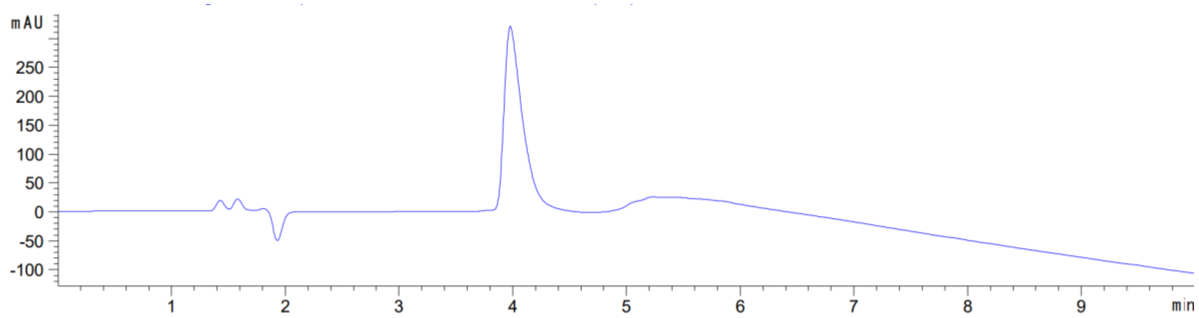


**H4R3me1**

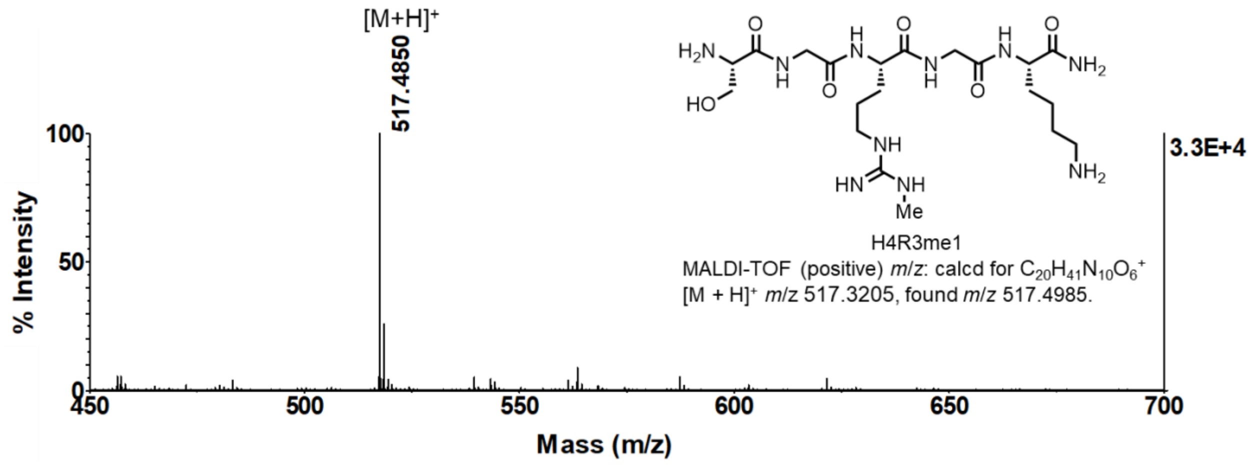


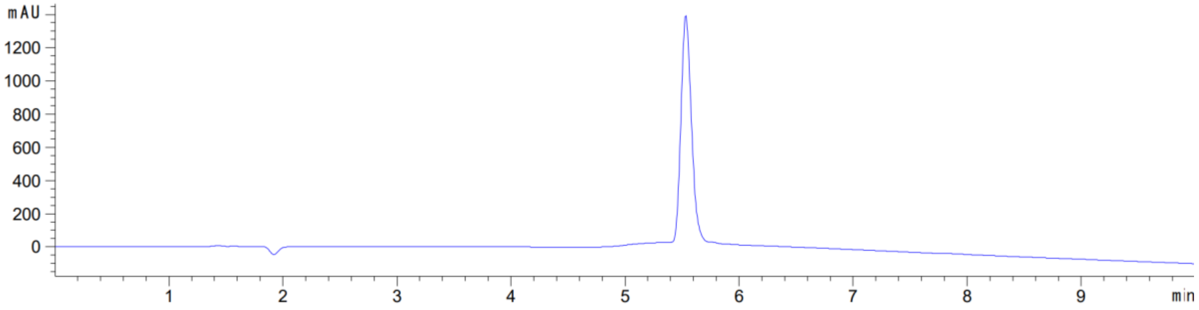

**H4R3me2a**

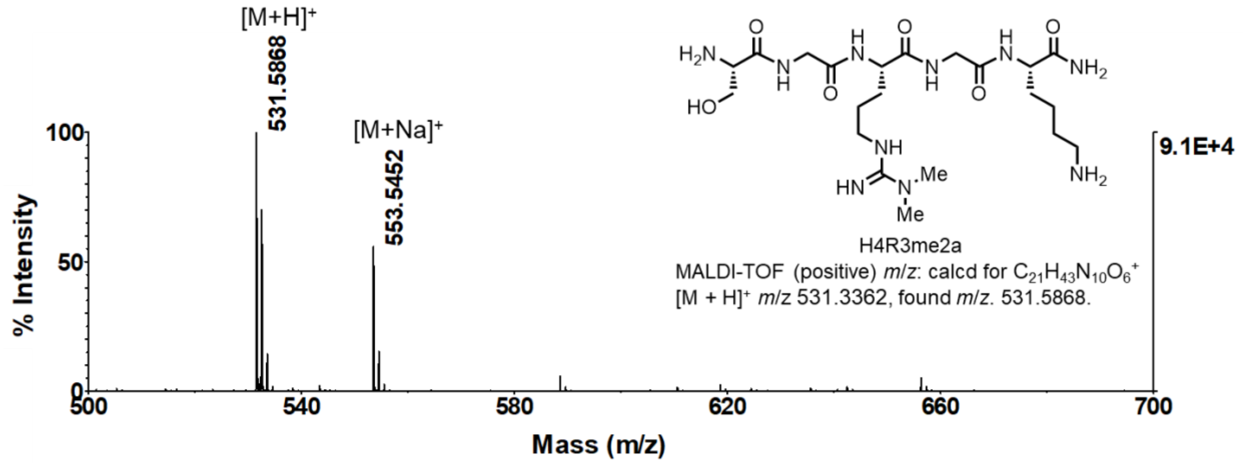


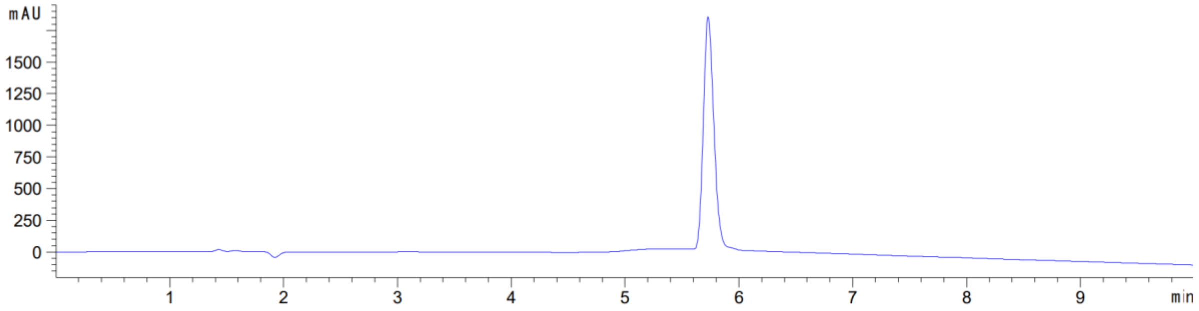


**H4R3me2s**

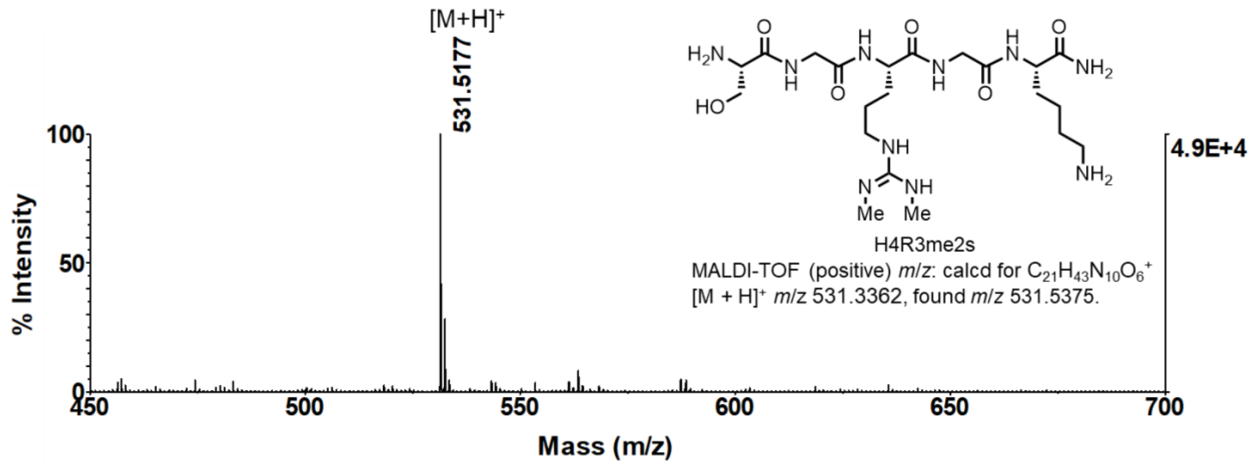


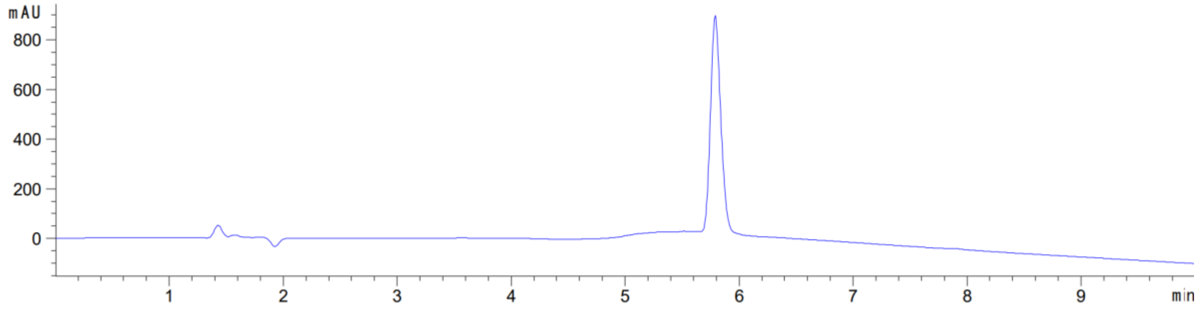

**H4K5ac**

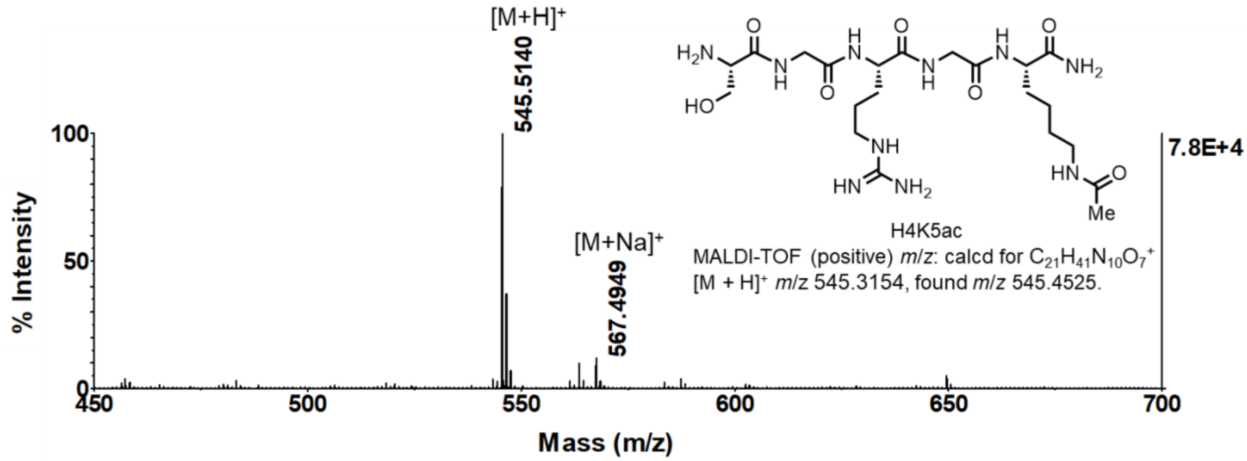
[M+H]+
545.5140
[M+Na]+
567.4949
100
50
0
% Intensity
450
500
550
600
650
700
Mass (m/z)
7.8E+4
H4K5ac
MALDI-TOF (positive) m/z: calcd for $C_{21}H_{41}N_{10}O_7^+$
[M + H]+ m/z 545.3154, found m/z 545.4525.

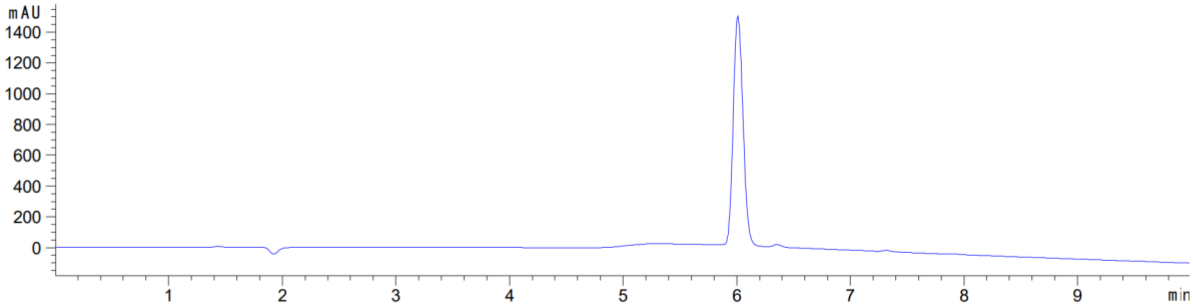
mAU
1400
1200
1000
800
600
400
200
0
1
2
3
4
5
6
7
8
9
min
